# Molecular mechanisms behind the functional (non)redundancy of CK1 paralogs in the Wnt pathway

**DOI:** 10.64898/2026.09.08.750032

**Authors:** Tomáš Gybeĺ, Kristína Gömöryová, Jana Bartošíková, Tereza Číhalová, Sara Bologna, Miroslav Micka, Ganji Sri Ranjani, David Potěšil, Ondrej Šedo, Tomasz W. Radaszkiewicz, Zbyněk Zdráhal, Konstantinos Tripsianes, Gopal P. Sapkota, Vítězslav Bryja

**Affiliations:** Department of Experimental Biology, Faculty of Science, Masaryk University, Brno, 62500, Czech Republic; CEITEC-Central European Institute of Technology, Masaryk University, Brno, 62500, Czech Republic; Medical Research Council Protein Phosphorylation & Ubiquitylation Unit, School of Life Sciences, University of Dundee, Dundee DD1 5EH, UK

**Author notes:** Correspondence to: prof. Mgr. Vítězslav Bryja, Ph.D., Department of Experimental Biology Faculty of Science, Masaryk University Kamenice 5, D36/112, 625 00 Brno Czech Republic.

**Keywords:** casein kinase 1 alpha (CK1α), casein kinase 1 epsilon (CK1ε), Wnt/β-catenin pathway, redundancy, protein chimera, Wnt signalosome, β-catenin destruction complex (degradasome), SACK1 (FAM83)

## Abstract

The casein kinase 1 (CK1) family of serine/threonine protein kinases consists of seven isoforms in humans. CK1 family members are important regulators of the Wnt/β-catenin signaling pathway. Using a comprehensive panel of CRISPR/Cas9-generated knockout cell lines, we demonstrated the opposing roles of endogenous CK1α (negative, via phosphorylation of β-catenin in the destruction complex) and CK1δ/ε (positive, via phosphorylation of DVL in the signalosome), while no phenotype was observed for CK1γ1/2/3 triple knockout cells. Using *in vitro* kinase assays and TurboID-based interactomics, we revealed that this functional divergence between CK1α and CK1ε is not due to the intrinsically different capacity to phosphorylate β-catenin or DVL but rather due to different affinities for the destruction complex and signalosome in the cellular environment. Through functional analysis of CK1α–CK1ε chimeras containing both N-terminal and C-terminal domain swaps, we identified the N-terminal lobe of CK1α and the C-terminus of CK1ε as determinants mediating increased affinity towards the degradasome and signalosome, respectively. We further show that the CK1α N-lobe not only drives cellular activity toward β-catenin but also underlies the CK1α-specific interaction with the scaffolding protein SACK1G (also known as FAM83G and PAWS1). Additionally, despite clearly distinct physiological roles of CK1α and CK1δ/ε, we provide evidence that, in the absence of CK1α, CK1δ and CK1ε can physically and functionally substitute for CK1α in the β-catenin destruction complex. This rewires CK1δ and CK1ε as negative regulators acting via phosphorylation of β-catenin in the destruction complex. These findings resolve prior contradictions by (i) clarifying context-dependent CK1 roles, (ii) identifying mechanistic determinants navigating CK1α and CK1ε to different substrates, and (iii) defining the limitations of these affinity-based subcellular distributions that become apparent especially in the physical absence of the physiological, high-affinity kinase. An important implication of our findings is the identification of a mechanism that changes the ultimate outcome of CK1δ/ε inhibitor treatment from Wnt/β-catenin pathway inhibition to its robust activation.

## INTRODUCTION

The canonical Wnt/β-catenin signaling pathway is a fundamental cellular communication system, conserved throughout evolution, that plays a pivotal role in embryonic development, tissue homeostasis, and stem cell regulation (Holzem et al., 2024; Steinhart and Angers, 2018). Its dysregulation is a hallmark of numerous human diseases, most notably cancer (Xue et al., 2025; Zhan et al., 2017). The pathway’s core mechanism revolves around the tightly controlled stabilization of the transcriptional co-activator β-catenin. In the absence of a Wnt ligand, a multiprotein β-catenin destruction complex—also termed as “degradasome”—orchestrated by the scaffold proteins Axin and APC, facilitates the sequential phosphorylation of β-catenin. This phosphorylation, initiated by Casein Kinase 1 (CK1) and followed by Glycogen Synthase Kinase 3 (GSK3), marks β-catenin for ubiquitination and subsequent proteasomal degradation, thus keeping the pathway inactive. The binding of a Wnt ligand to its Frizzled and LRP5/6 co-receptors triggers recruitment of the destruction complex to the plasma membrane, inhibiting its activity and allowing newly synthesized β-catenin to accumulate, enter the nucleus, and activate target gene expression (Gammons and Bienz, 2018; Maurice and Angers, 2025).

Within this critical regulatory network, the CK1 family of serine/threonine kinases has long been recognized as a central player (Vielhaber and Virshup, 2001). However, its precise role is paradoxically complex and full of ambiguities (Polakis, 2002). The family comprises seven highly homologous isoforms/paralogs in humans (α, α-like, γ1, γ2, γ3, δ, ε), encoded by distinct genes (Fulcher and Sapkota, 2020). Phosphorylation by CK1 has been reported for almost every central protein of the Wnt pathway: FZD and LRP5/6 receptors, the key signaling intermediate Dishevelled, and destruction complex proteins such as Axin, APC and β-catenin itself (Cruciat, 2014; Maurice and Angers, 2025). In line with these observations, CK1 kinases have been functionally implicated at nearly every level of the Wnt pathway.

Despite some conflicting observations, the data synergize in the prevailing model which, however, presents a stark functional dichotomy for individual CK1 paralogs: CK1α is considered a key negative regulator of Wnt/β-catenin signaling, acting via the priming phosphorylation on β-catenin required for subsequent phosphorylation by GSK3 and β-catenin degradation (Polakis, 2002; Stamos and Weis, 2013). In contrast, CK1δ and CK1ε are viewed as positive regulators, acting at the level of the plasma membrane signalosome to phosphorylate components such as Dishevelled (DVL) and the LRP5/6 co-receptor, thereby promoting pathway activation (Bryja et al., 2007; Cruciat, 2014; Dale, 2006). The molecular basis by which these highly similar kinases achieve diametrically opposed functions remains a major unresolved question. For instance, CK1α shares higher overall sequence homology (**Fig.** 1A) with the functionally inert CK1α-like isoform than the positive regulators CK1δ and CK1ε share with each other, despite the latter pair exhibiting clear functional redundancy (Bryja et al., 2007; Gybeľ et al., 2024; Morgenstern et al., 2017). This suggests that features beyond the conserved parts of the kinase domain must dictate their specific roles and substrate choices. Whether this specificity arises from inherent substrate selectivity, subcellular localization, or interaction with distinct regulatory partners is not well understood. Furthermore, the contribution of the less-characterized amino- (N-) and carboxyl- (C-) terminal regions, which are the most divergent parts among isoforms, has been largely overlooked.

**Figure 1:**
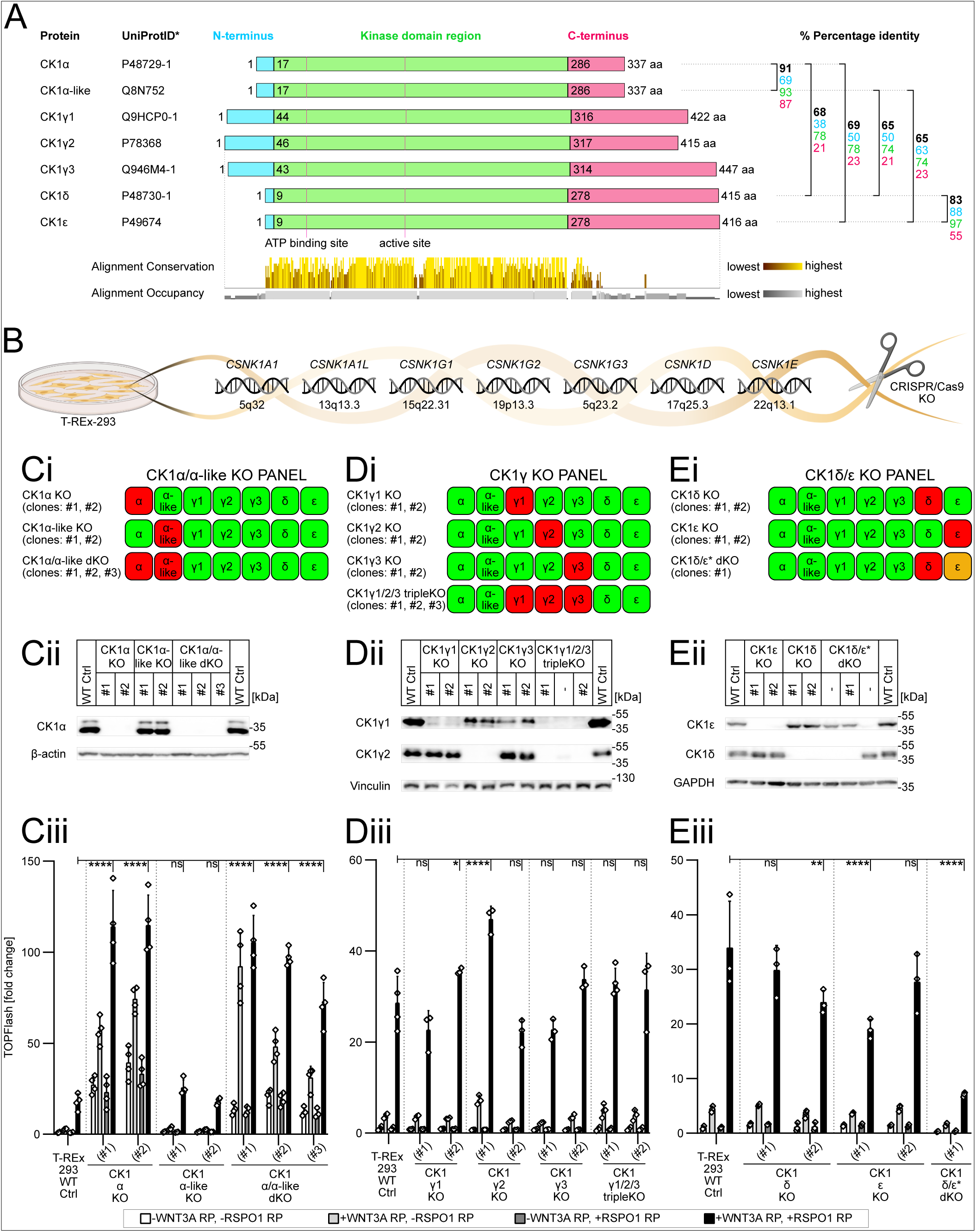
CK1α and CK1δ/ε exhibit opposite roles in the Wnt/β-catenin signaling despite their high homology. **(A)** Schematic depiction of the seven isoforms of the CK1 family. The isoforms share high homology in their kinase domains (shown in green) but differ in their N-terminal (blue) and C-terminal (red) regions. The alignment was produced by ClustalOWS and, together with the conservation plot, visualized using Jalview 2.11.4.1. The percentage identity is based on pairwise alignment using BLOSUM62 from the ClustalOWS alignment (see Fig.S1A-D), presented as follows: the overall alignment is shown in black; alignments of the N-termini, kinase domains, and C-termini are shown in their respective color codes. *Canonical variants. **(B)** CK1 isoforms are encoded by seven genes located at different chromosomal positions in the human genome (GRCh38/hg38 genome assembly). The depicted genes were knocked out in T-REx-293 cells using CRISPR/Cas9 editing. **(Ci/Di/Ei)** Schematic depiction of CK1α/α-like KO panel (Ci), CK1γ KO panel (Di), CK1δ/ε KO panel (Ei), along with the KO cell lines prepared in these panels. Red = KO, orange = partial KO. **(Cii/Dii/Eii)** Western blot validation of single and compound KO lines from the corresponding KO panels. **(Ciii/Diii/Eiii)** TOPFlash analysis of CK1 isoform KO panel lines treated for 16 hours with combinations of Wnt-3a (WNT3A) recombinant protein (80 ng/ml) and/or R-spondin-1 (RSPO1) recombinant protein (25 ng/ml). All treatments included LGK974 (0.5 µM). Normalized ratios of TOPFlash luciferase to Renilla luciferase relative luminescence units are plotted (fold change). Statistical analysis: Ordinary two-way ANOVA with Tukey’s multiple comparisons test (ns [P>0.05], * [P≤0.05], ** [P≤0.01], *** [P≤0.001], **** [P≤0.0001]). Columns represent means with S.D. error bars. n = 3-4 independent biological replicates. Figures Cii and Ciii reproduce data previously presented in Gybel et al., 2024 (Fig. 3B/C).

In this study, we systematically deconstruct the functional specialization of CK1 isoforms to resolve these ambiguities. Using a comprehensive panel of CRISPR/Cas9-generated knockout cell lines, we first confirm the opposing roles of endogenous CK1α (negative) and CK1δ/ε (positive) in the Wnt/β-catenin pathway, while demonstrating that loss of all three CK1γ paralogs produces no phenotype in this model. We reveal that this functional divergence is not due to intrinsic differences in enzymatic activity but rather to preferential interactions within the components of either the degradasome or the signalosome. Analysis of CK1α and CK1ε chimeras pinpointed the structural determinants of this specificity to the N-terminal lobe of CK1α for the degradasome, and to the C-terminal tail of CK1ε for the signalosome. Interestingly, the region that mediates cellular association with the degradasome is both sufficient and required for interaction with the CK1 scaffolding protein SACK1G (also known as FAM83G and PAWS1). Despite these unique affinity determinants of CK1α and CK1ε physiological function, we uncover a context-dependent functional plasticity wherein CK1δ and CK1ε can be rewired to substitute for CK1α in the destruction complex, and as a consequence, can completely switch from positive to negative regulators of the Wnt/β-catenin pathway.

## RESULTS

### CK1α and CK1δ/ε exhibit opposing roles in the Wnt/β-catenin signaling despite their high homology

We decided to elaborate on our previous study (Gybeľ et al., 2024), which focused on CK1α, its splice variants, and CK1α-like isoform—their functions and redundancy in the canonical Wnt signaling pathway. Among the CK1 family members, isoforms CK1α and CK1α-like exhibit the highest overall homology in terms of amino acid sequence (**Fig.** 1A, see “% Percentage identity” black numbers; **Fig.** S1A). Despite this, we observed no substantial redundancy between CK1α and CK1α-like, nor activity of CK1α-like in the canonical Wnt pathway. On the other hand, CK1δ and CK1ε exhibit apparent redundant nature in the Wnt signaling (Bryja et al., 2007; Morgenstern et al., 2017). While their overall homology is lower than that between CK1α and CK1α-like, their homology within the kinase domain region is higher (**Fig.** 1A and **Fig.** S1C).

To comprehensively address the issue of functional requirement and redundancy of CK1 proteins, we produced a series of CK1-deficient cells using CRISPR/Cas9 (**Fig.** 1B). We generated three panels of CK1 knockout (KO) lines in T-REx-293 cells: CK1α/α-like KO panel (**Fig.** 1Ci) (originally published in (Gybeľ et al., 2024)), CK1γ KO panel (**Fig.** 1Di), and CK1δ/ε KO panel (**Fig.** 1Ei). At least two clones for each KO type were isolated, and monoclonal cell lines were established. Characterization of the generated cell lines by restriction fragment length polymorphism, next-generation sequencing, and Western blotting (WB) is provided in **Fig.** 1Cii/Dii/Eii, **Fig.** S2 and **Fig.** S3.

The CK1 KO panels were further tested for their ability to transduce Wnt/β-catenin signaling. Cells were pretreated with the porcupine inhibitor LGK974 to prevent Wnt ligand secretion, and pathway activity was examined under four conditions: (i) no stimulation, (ii) stimulation with the prototypical canonical Wnt ligand—Wnt3a, (iii) stimulation with the potentiator of canonical Wnt signaling—R-spondin-1, and (iv) combined Wnt3a and R-spondin-1 treatment for maximal pathway stimulation. The parental T-REx-293 wild-type (WT) cells served as controls (WT Ctrl). TOPFlash assays monitoring TCF/LEF-dependent transcription (**Fig.** 1Ciii/Diii/Eiii) and WB analyses of Wnt pathway proteins served as primary readouts.

CK1α/α-like double KO (dKO) cells (**Fig.** 1Ciii) exhibited significantly higher Wnt/β-catenin pathway activity, consistent with the known role of CK1α in the destruction complex (Liu et al., 2002). As we showed previously, this activity is unique to CK1α and not CK1α-like (Gybeľ et al., 2024). CK1γ KO cells (**Fig.** 1Diii and **Fig.** S2E) showed no significant alterations in the Wnt/β-catenin signaling that would be consistent across clones. While we successfully generated single KOs of CK1δ and CK1ε without complications, repeated attempts to isolate CK1δ/ε dKO lines failed, likely due to the reported synthetic lethality of CK1δ and CK1ε isoforms (Morgenstern et al., 2017). Surprisingly, we isolated one clone, designated CK1δ/ε* dKO (#1), which is a full KO for CK1δ and a compound mutant for CK1ε: one allele KO, the other with a 3 bp in-frame insertion plus a 2 bp change resulting in a CK1ε M^58^M^59^ to N^58^L^59^P^60^ mutant. This mutation produces a CK1ε hypomorph with reduced activity, as shown by functional assays (**Fig.** S3D/E). CK1δ/ε* dKO cells demonstrated partial defects in Wnt/β-catenin signaling, evidenced by lower Wnt3a-induced TOPFlash response (**Fig.** 1Eiii) and absence of Wnt-3a-induced phosphorylation-dependent mobility shift of DVL3 (**Fig.** S3F). However, these cells still responded to Wnt3a via LRP6 phosphorylation, Axin1 dephosphorylation, and increased active (dephosphorylated) β-catenin levels (**Fig.** S3F).

In conclusion, our data from CK1 KO T-REx-293 lines confirm the predominantly accepted view that CK1 paralogs have clearly distinct and non-redundant functions in the Wnt/β-catenin pathway. More specifically, CK1α functions as a negative regulator, while CK1δ and CK1ε act as positive regulators of the pathway. We observed no functional phenotype following the depletion of CK1γ.

### CK1 isoforms exhibit preferential interactors *in vivo* without substrate selectivity *in vitro*

To get a better insight into the differences in the biological functions of CK1 paralogs, we decided to explore the proximity interactome of CK1 paralogs using TurboID of CK1α (splice variant v2), CK1α-like, CK1δ, and CK1ε (**Fig.** 2A). We prepared Flp-In T-REx-293 stable cell lines with TetON-inducible expression of N-terminally TurboID-tagged CK1s; TurboID-only was used as a negative control, whereas TurboID-DVL3 served as a positive control for signalosome interactors. Proximity biotinylation was triggered by biotin supplementation, biotinylated proteins were pulled down using streptavidin beads, and then analyzed by mass spectrometry. The enrichment of individual preys—i.e., endogenous proteins in the proximity of TurboID-tagged baits—over TurboID alone was quantified using fold change and adjusted p-value (see Materials and Methods for details). Surprisingly, we detected key components of both destruction complex and the signalosome interacting with all analyzed CK1 kinases (**Fig.** 2B). However, CK1α (and in some cases CK1α-like) interacted much more strongly with the proteins forming the degradasome–i.e., β-catenin, APC1/2 and Axin1/2—whereas CK1δ and CK1ε interacted preferentially with the proteins forming the signalosome–i.e., FZD receptors, DVL1/2/3 and LRP5/6. This suggests that there is no strict exclusivity in the interaction between individual CK1 kinases and particular preys; rather, the cellular outcome is determined by differences in affinity–relatively higher for CK1α and degradasome proteins, and for CK1δ and CK1ε with signalosome proteins.

**Figure 2:**
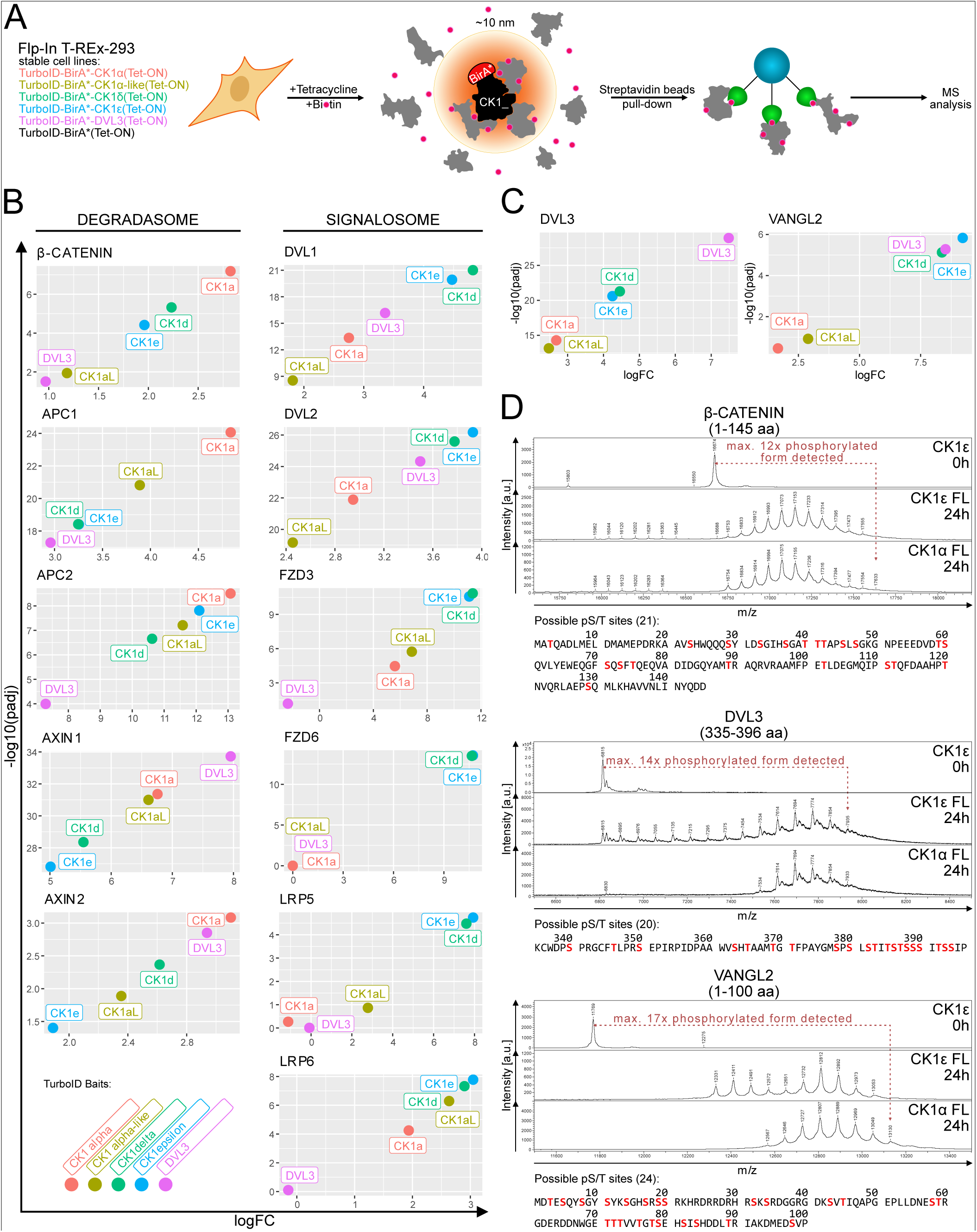
CK1 isoforms exhibit preferential interactors *in vivo* without substrate selectivity *in vitro*. **(A)** Scheme of the TurboID experiment. Individual tetracycline-inducible TurboID-BirA*-CK1 isoform cell lines prepared in Flp-In T-REx-293 cells, were induced with tetracycline overnight and biotin for 30 minutes. All CK1 isoform-interacting proteins within an approximate 10 nm radius were biotinylated. These proteins were precipitated with streptavidin beads and submited for mass spectrometry analysis. **(B)** TurboID interactome analysis of CK1 isoforms and DVL3. Detected Wnt signaling-related hits (preys) are visualized by their intensities detected for individual TurboID-CK1/DVL baits. **(C)** Extension of Fig.2B showing hits (preys) DVL3 and VANGL2. **(D)** Matrix-assisted laser desorption/ionization-time of flight mass spectrometry (MALDI-TOF MS) analysis of three CK1 isoform substrates: fragments of β-catenin (1-145 aa), DVL3 (335-396 aa) and VANGL2 (1-100 aa).

In parallel, we compared the kinase activity of CK1α and CK1ε using *in vitro* kinase assays. We purified representative kinases: full-length (FL) CK1α and CK1ε (**Fig.** S4A). Additionally, we purified well-described interactors (**Fig.** 2C) and substrates of CK1 in the Wnt pathway: β-catenin (CK1α degradasome substrate), DVL3 (CK1δ/ε signalosome substrate), and Vangl2 (non-canonical Wnt pathway CK1δ/ε substrate) (**Fig.** S4B). *In vitro* kinase assay products were analyzed by Matrix-assisted laser desorption/ionization-time of flight mass spectrometry analysis (MALDI-TOF MS). MALDI intact mass analysis shows the phosphorylation status of the substrate as an increment in its molecular weight through characteristic 80 Da mass shifts that correspond to phosphate group additions. Interestingly, we did not observe any significant differences between CK1α and CK1ε on any of the studied substrates (**Fig.** 2D).

In summary, these results suggest that the observed differences in biological function *in vivo* are not driven by intrinsic substrate selectivity of CK1α or CK1ε, but rather by their affinities towards the degradasome and signalosome, respectively, within the complex cellular environment.

The N-lobe of CK1α and the C-terminus of CK1ε define cellular preferences for the β-catenin destruction complex and DVL3 signalosome, respectively.

To identify which region underlies the cellular preferences and functions of CK1α and CK1ε, we tested a series of chimeric CK1 proteins composed of regions from CK1α and CK1ε. Based on the alignment of the N-terminal (**Fig.** 3Ai) and C-terminal (**Fig.** 3Aii) regions of CK1α, CK1ε and CK1δ, we established a panel of mutant constructs (**Fig.** Aiii and **Fig.** 3B). We swapped both the N- and C-terminal regions of CK1α and CK1ε. Additionally, we produced the short (S) and long (L) truncations of the C-terminal regions.

**Figure 3:**
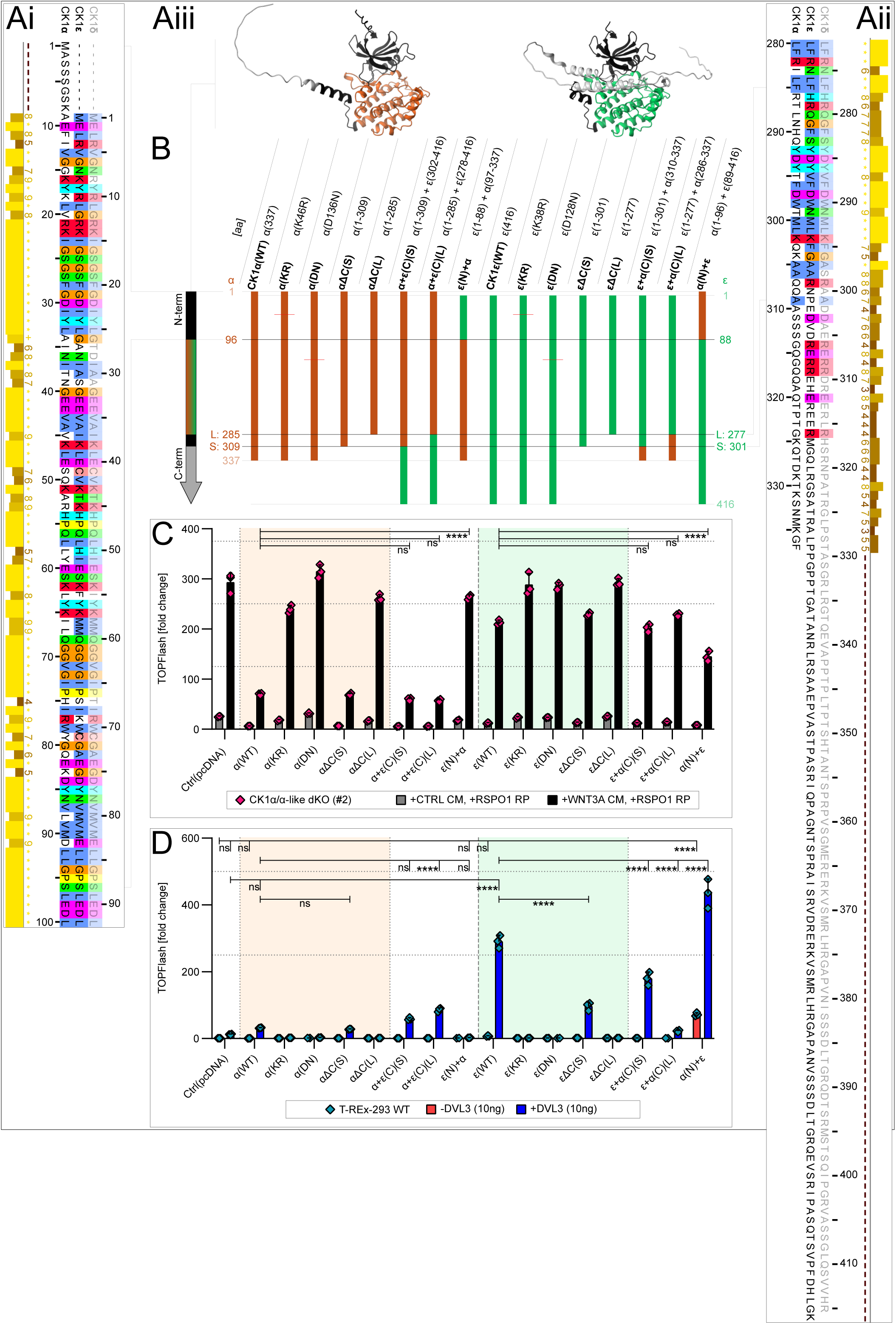
The N-lobe of CK1α and the C-terminus of CK1ε define cellular preferences for the β-catenin destruction complex and DVL3 signalosome, respectively. **(A)** Protein sequence alignment of the N-terminal regions (Ai) and C-terminal regions (Aii) of CK1α, CK1ε, and CK1δ, highlighting residues with higher variability (created in Jalview 2.11.4.1). AlphaFold3 structural predictions (Aiii) of CK1α (orange) and CK1ε (green) depict the localization of the N-terminal and C-terminal regions selected for swapping. **(B)** Schematic depiction of the panel showing N-terminal and C-terminal single swapping mutants between CK1α and CK1ε. **(C)** TOPFlash assay: degradasome assay with mutants from Fig.3B. Statistical analysis: Three-way ANOVA with Tukey’s multiple comparisons test. **(D)** TOPFlash assay: signalosome assay with mutants from Fig.3B. Statistical analysis: Ordinary two-way ANOVA with Tukey’s multiple comparisons test. **(C,D)** Columns represent means with S.D. error bars. n = 3 independent biological replicates. (ns [P>0.05], * [P≤0.05], ** [P≤0.01], *** [P≤0.001], **** [P≤0.0001]).

To investigate the functional impact of these mutants on the Wnt signaling pathway, we conducted two distinct assays assessing the negative role of CK1 in the destruction complex (degradasome assay) and the positive role of CK1 associated with DVL phosphorylation (signalosome assay). Both assays are based on the TOPFlash reporter. The degradasome assay, introduced previously (Gybeľ et al., 2024), measures the rescue of hyperactivated Wnt signaling in CK1α/α-like dKO cells back to normal levels. The signalosome assay is based on low-level transfection of DVL3 and/or CK1ε into T-REx-293 WT cells. Transfection of DVL3 or CK1ε alone fails to induce a signaling response, whereas co-transfection results in robust, synergistic upregulation of TOPFlash.

The analysis of all CK1 chimeras in both assays is shown in **Fig.** 3C (degradasome assay) and **Fig.** 3D (signalosome assay). Expression levels of individual CK1 mutants and chimeras are shown in **Fig.** S5B/C. Among the 16 CK1 constructs tested, only those with an intact and active CK1α kinase domain showed full activity in the degradasome assay (**Fig.** 3C). CK1α activity was lost by mutations interfering with kinase activity (K-R, D-N) or by a large deletion of the C-terminus ΔC(L), which we believe also affected kinase activity. The shorter deletion of the C-terminus (αΔC(S)) or chimeras containing CK1ε C-termini (α+ ε(C)) in both lengths, S or L, did not affect activity in the degradasome. In contrast, swapping the N-terminal region of CK1α for that of CK1ε (ε(N)+α) fully compromised CK1α’s ability to function in the degradasome. CK1ε showed only mild rescue, which was enhanced in the chimera containing the N-terminus from CK1α (α(N)+ε). We conclude that the N-lobe of CK1α is the critical determinant—both necessary and sufficient— to mediate CK1α’s cellular activity in the destruction complex.

In the signalosome assay (**Fig.** 3D), CK1ε was, as expected, much more active than CK1α. Similarly to CK1α in the degradasome assay, CK1ε activity in the signalosome assay was lost by mutations interfering with kinase activity (K-R, D-N) or by a larger deletion of the C-terminus ΔC(L), presumably interfering with kinase activity as well. CK1ε activity was also significantly diminished by truncation of its C-terminal region (εΔC(S)) or its replacement with the CK1α C-terminus (ε+α(C)(S/L)). Conversely, CK1α’s poor performance in the signalosome assay was improved by addition of the CK1ε C-terminus (see α+ ε(C)(S/L)) in both lengths, S or L. These data collectively suggest that the CK1ε C-terminus mediates CK1 activity towards DVL and the signalosome.

A striking phenotype was observed in the chimera composed of the N-lobe of CK1α and the rest of CK1ε (α(N)+ε). This chimera was hyperactive, capable of triggering the signaling even in the absence of co-expressed DVL3 (red bar), and surpassed CK1ε(WT) activity in the presence of DVL3 (blue bar) (**Fig.** 3D). This phenotype was confirmed by WB analysis (**Fig.** S5C), where this mutant caused the most prominent hyperphosphorylation of DVL3. The hyperactivity of this chimera depends on both its kinase function (**Fig.** S5D and S5E) and an intact C-terminal region of CK1ε (**Fig.** S7C). The precise mechanism is unclear, but one plausible explanation is that the structural elements mediating interactions with the degradasome (CK1α N-lobe) and signalosome (CK1ε C-terminus) in one protein recruit the destruction complex to the membrane, where it becomes inactivated.

We generated two additional series of CK1 chimeras. First, smaller swaps of CK1α-specific N-lobe segments (**Fig.** S6) produced partial phenotypes, suggesting that the unique N-lobe surface consists of discontinuous motifs. Second, we created double swapping mutants where the kinase core belongs to one isoform while the N- and C-terminal regions belong to the other (**Fig.** S7). However, these did not yield more profound effects on phenotype rescue. On contrary, the larger the portion of CK1α in the C-terminus of CK1ε, the poorer its performance in the signalosome assay, as observed in single (**Fig.** 3D) and double swapping mutants (**Fig.** S7C). A similar trend was visible to some extent also in the degradasome assay (**Fig.** 3C and **Fig.** S7B).

Based on this extensive set of experiments (**Fig.** 3 and **Fig.** S5-S7), we conclude that the N-lobe of CK1α and the C-terminus of CK1ε are the key determinants controlling the selective activity of CK1α in the degradasome and CK1ε in the signalosome, respectively.

The N-terminal region of CK1α is a structural determinant of its selective interaction with SACK1G (FAM83G).

One possible mechanism for the functional specificity of individual CK1 paralogs is via their interaction with <u>S</u>caffold <u>A</u>nchor of <u>CK1</u> (SACK1)-domain-containing proteins A-H (SACK1A-H), formerly known as the family of sequence similarity 83 (FAM83). CK1α has been reported to interact with all eight (A-H) members of the SACK1 family, including SACK1G (FAM83G), while CK1ε associates only with SACK1A/B/E/H (FAM83A/B/E/H) (Fulcher et al., 2018). Our TurboID assay confirmed this interaction profile, showing CK1α in complex with all three endogenously expressed SACK1 proteins—SACK1B, SACK1G and SACK1H— whereas CK1δ/ε clearly failed to interact with SACK1G (**Fig.** 4A).

**Figure 4:**
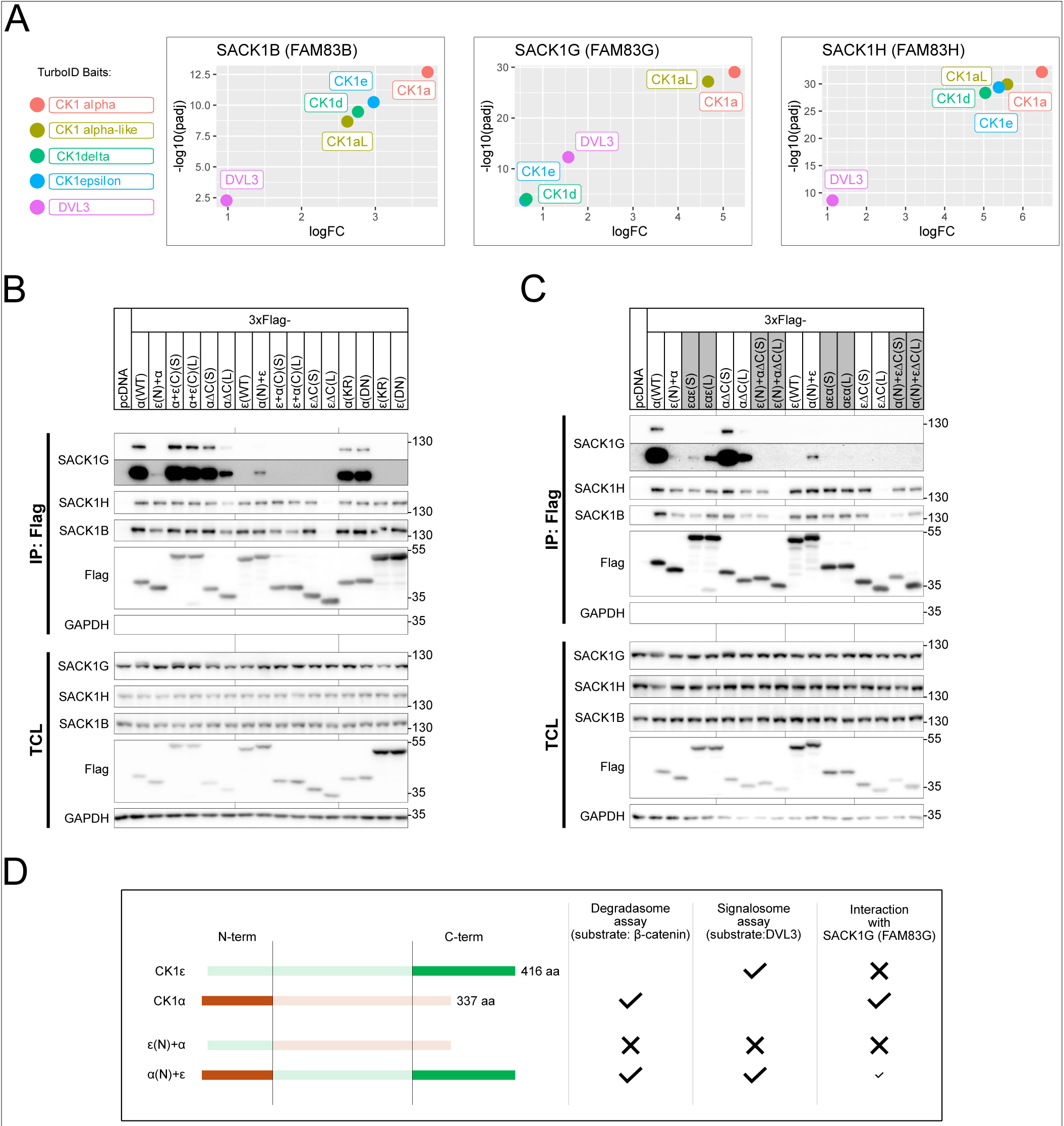
The N-terminal region of CK1α is a structural determinant of its selective interaction with SACK1G (FAM83G). **(A)** TurboID interactome analysis of CK1 isoforms and DVL3. Detected Wnt signaling-related hits (preys: SACK1B, SACK1G, SACK1H) are visualized by their intensities detected for individual TurboID-CK1/DVL baits. **(B)** Co-immunoprecipitation: Immunoprecipitation of 3xFlag-tagged CK1 isoforms, their mutants, and chimeric constructs (panel from Fig.3B) in overexpression conditions, with detection of pulled-down SACK1G, SACK1H, and SACK1B. Three biological replicates are presented (Fig.S8A). TCL - total cell lysate input **(C)** Co-immunoprecipitation: Immunoprecipitation of 3xFlag-tagged CK1 isoforms, their mutants, and chimeric constructs (panel from Fig.S7A) in overexpression conditions, with detection of pulled-down SACK1G, SACK1H, and SACK1B. Three biological replicates are presented (Fig.S8B). TCL - total cell lysate input **(D)** Graphical summary of Fig.3 and Fig.4.

Importantly, the interaction between CK1α and SACK1G has been shown to be essential for proper Wnt/β-catenin signaling function (Bozatzi et al., 2018; Glennie et al., 2024). To test whether the function of our CK1 chimeras correlates with their capacity to interact with SACK1G, we performed a set of co-immunoprecipitation experiments (**Fig.** 4B/C and **Fig.** S8A/B). SACK1G interacted clearly only with CK1α and not with CK1ε, consistent with previous reports (Fulcher et al., 2018). Interestingly, the CK1ε(N)+α chimera, which exhibited impaired function in the degradasome assay, also lost the ability to interact with SACK1G. This suggests that the N-terminal lobe of CK1α mediates the specific interaction with SACK1G. This motif is not only required but also sufficient, as demonstrated by the gain of interaction between the CK1α(N)+ε chimera and SACK1G. However, the N-terminal lobe of CK1α alone is not sufficient to establish interaction with SACK1G, as shown in the CK1α splice variant v9 (**Fig.** S8C/D). This indicates that other structural elements, shared between CK1α and CK1ε, are required to support this interaction.

In summary (**Fig.** 4D), the activity of CK1 variants in the degradasome assay correlates with their ability to interact with SACK1G through the N-terminus of CK1α. We propose that the specific interaction of CK1α with SACK1G contributes to its proper subcellular localization and affinity for the destruction complex, thereby mediating its negative regulatory role in the Wnt/β-catenin signaling.

### In the absence of CK1α, CK1δ and CK1ε physically and functionally replace CK1α in the destruction complex

Our data–especially from the TurboID dataset (**Fig.** 2B/C) and the degradasome assay (**Fig.** 5A and **Fig.** S9A)–suggest that despite clear structural determinants mediating affinity to the degradasome or signalosome, CK1δ and CK1ε can at least partially replace CK1α in the destruction complex. We also show that in the presence of physiological levels of CK1α, this replacement is unlikely; but what happens in the absence of CK1α? We decided to test this hypothesis through a series of experiments.

**Figure 5:**
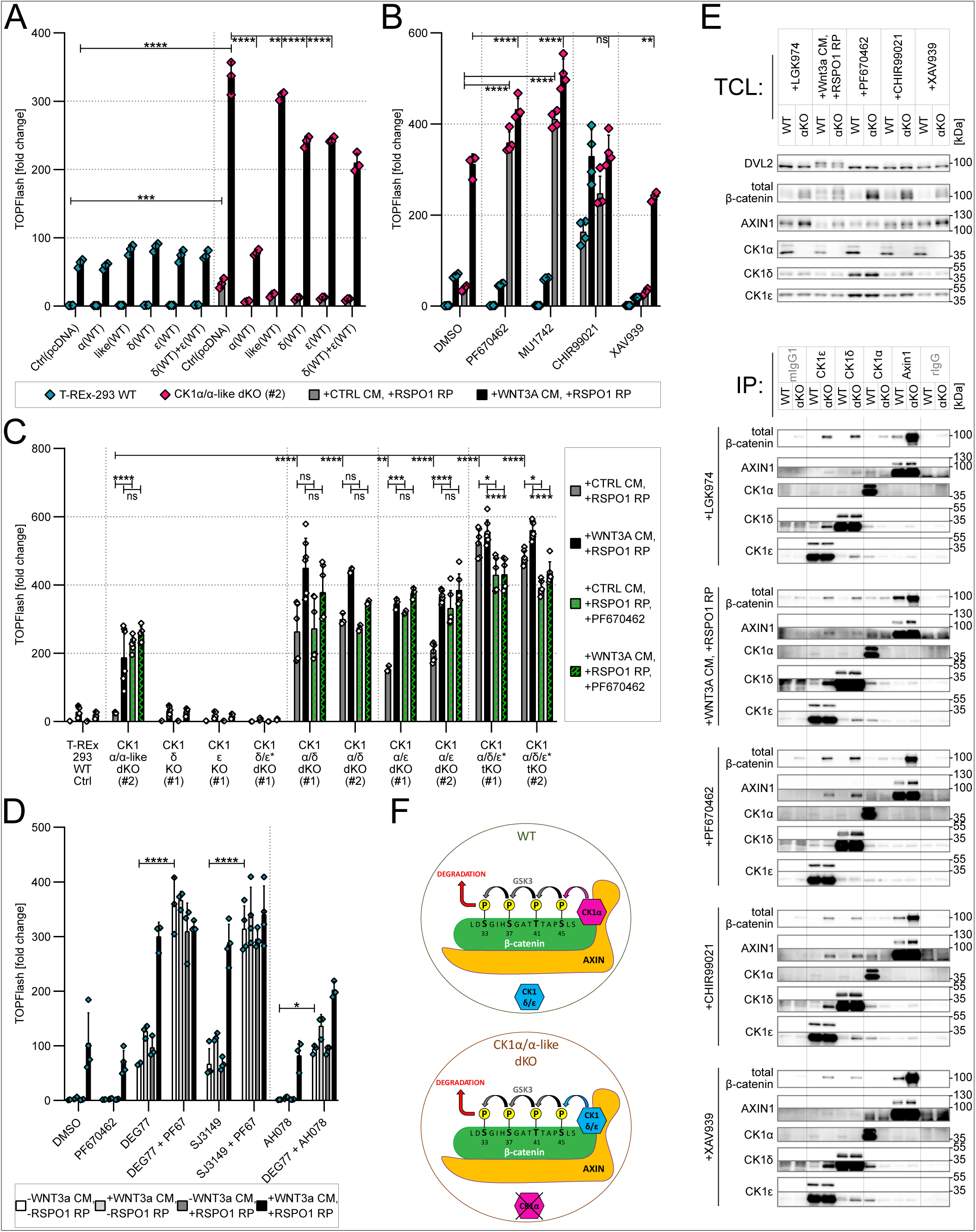
In the absence of CK1α, CK1δ and CK1ε physically and functionally replace CK1α in the destruction complex. **(A)** TOPFlash assay: degradasome assay with CK1 isoforms, showing their potential to act as negative regulators of the Wnt/β-catenin pathway. Statistical analysis: Three-way ANOVA with Tukey’s multiple comparisons test. n = 3 independent biological replicates. **(B)** TOPFlash assay testing pharmacological inhibition of Wnt/β-catenin pathway components using PF670462 (CK1δ/ε inhibitor; 10 µM), MU1742 (CK1α/δ/ε inhibitor; 10 µM), CHIR99021 (GSK3α/β inhibitor; 5 µM) and XAV939 (TNKS1/2 inhibitor; 10 µM) in WT and CK1α/α-like dKO cells. Treatments were carried out for 16 hours along with Wnt/β-catenin pathway stimulation in the presence of 0.5 µM LGK974. Statistical analysis: Three-way ANOVA with Tukey’s multiple comparisons test. n = 4 independent biological replicates. **(C)** TOPFlash assay testing genetic models of CK1α/δ/ε single and compound depletion in the absence or presence of PF670462 (10 µM/16 h) and Wnt/β-catenin pathway stimulation in the presence of 0.5 µM LGK974. Statistical analysis: Three-way ANOVA with Tukey’s multiple comparisons test. n = 3-7 independent biological replicates. **(D)** TOPFlash assay testing pharmacological inhibition and/or depletion of CK1α/δ/ε isoforms using PF670462 (CK1δ/ε inhibitor; 10 µM), DEG77 (1 µM), SJ3149 (1 µM) and AH078 (5 µM) in WT cells. Treatments were carried out for 16 hours along with Wnt/β-catenin pathway stimulation in the presence of 0.5 µM LGK974. Statistical analysis: Ordinary two-way ANOVA with Tukey’s multiple comparisons test. n = 3-4 independent biological replicates. **(A,B,C,D)** (ns [P>0.05], * [P≤0.05], ** [P≤0.01], *** [P≤0.001], **** [P≤0.0001]). Columns represent means with S.D. error bars. **(E)** Endogenous co-immunoprecipitations under Wnt-ON/OFF conditions. WT and CK1α/α-like dKO (#2; αKO) cells were treated for 16 hours with WNT3A CM and RSPO1 RP, PF670462 (10 µM), CHIR99021 (2 µM), XAV939 (10 µM), and 0.5 µM LGK974 in all conditions. **(F)** Suggested model of redundancy in which CK1δ/ε substitute CK1α in its absence within the β-catenin destruction complex.

First, is endogenous CK1δ/ε kinase activity required for the phenotype of CK1α KO cells, i.e., upregulated TOPFlash activity? Surprisingly, when we treated CK1α KO cells with CK1δ/ε-selective inhibitors (PF670462 and MU1742 (Němec et al., 2023)) (**Fig.** 5B and **Fig.** S9B/C), we observed robust and Wnt-3a-independent activation of TOPFlash. This suggests a substantial contribution of CK1δ/ε to the destruction complex function in the absence of CK1α.

We also explored other components of the destruction complex in the absence of CK1α using pharmacological inhibition. We tested CHIR99021—the inhibitor of glycogen synthase kinase 3 (GSK3)— and XAV939—the inhibitor of tankyrase (TNKS) 1 and 2 (**Fig.** 5B and **Fig.** S9B/C). Stabilization of Axin1, and thus the destruction complex, through TNKS1/2 inhibition suppressed signaling in both WT and CK1α KO cells, as expected. GSK3 inhibition increased basal signaling in both WT and CK1α KO cells. Interestingly, GSK3 inhibition activated the pathway only to levels that did not exceed those observed in CK1α KO cells treated with CK1δ/ε inhibitors. This suggests that the combination of CK1α depletion and CK1δ/ε inhibition leads to complete inhibition of the priming phosphorylation of β-catenin.

Because pharmacological inhibition of CK1δ/ε can lead to their accumulation and potential artifacts, we validated CK1δ/ε’s contribution to the destruction complex function using genetic models. Single and compound KO lines of CK1α, CK1δ and CK1ε were analyzed (**Fig.** 5C and **Fig.** S9D). The CK1α/δ, CK1α/ε, and CK1α/δ/ε* cells recapitulated the phenotype observed with pharmacological CK1δ/ε inhibition in CK1α KO cells (**Fig.** 5B)–i.e., Wnt-independent hyperactivation of the Wnt/β-catenin pathway. This finding supports the role of CK1δ/ε in the destruction complex of CK1α KO cells but is surprising because it suggests that in the absence of CK1α, deletion of either CK1δ or CK1ε is sufficient to further activate the pathway.

As CRISPR KO lines can exhibit some adaptation due to their production process, we confirmed the phenotype—the unexpected role of CK1δ/ε in CK1α absence—using state-of-the-art CK1α (SJ3149 and DEG77) (Nishiguchi et al., 2024; Park et al., 2023) and CK1δ/ε (AH078) (Haag et al., 2025) protein degraders. As shown in **Fig.** 5D and **Fig.** S9E, the TOPFlash assay with combinations of PF670462 and CK1α degraders in WT T-REx-293 cells repeatedly confirmed Wnt/β-catenin pathway upregulation by CK1δ/ε inhibitors only in the absence of CK1α.

Functional data suggest that CK1δ/ε physically replaces CK1α in the destruction complex once CK1α is removed genetically or pharmacologically. To confirm this experimentally, we performed endogenous co-immunoprecipitations in WT and CK1α KO cells under various states of Wnt signaling. As shown in **Fig.** 5E, Axin1, a key component of the destruction complex (van Kappel and Maurice, 2017), was found in complex with CK1δ and CK1ε, respectively, only in CK1α-deficient cells. This biochemically confirms that, in the absence of CK1α, CK1δ/ε physically bind Axin1 and can act at the degradasome (**Fig.** 5F).

Finally, to confirm that the rewiring of CK1δ/ε function in CK1α absence occurs downstream of the signalosome, we produced compound KOs lacking both CK1α and LRP5/6 or DVL1/2/3 (**Fig.** S9F). Lack of LRP5/6 or DVL1/2/3 causes cells to be completely unresponsive to Wnt-3a-induced signaling (Bernatik et al., 2021; Paclíková et al., 2017). However, CK1δ/ε inhibition by PF670462 in both CK1α+LRP5/6 and CK1α+DVL1/2/3 compound KO cells again led to robust activation of TOPFlash–similar to single CK1α KOs. This independently confirms that CK1δ/ε acquire a novel function downstream of the receptor complex– in the β-catenin destruction complex—in the absence of CK1α (**Fig.** S9G).

## DISCUSSION

The canonical Wnt/β-catenin pathway is governed by a series of phosphorylation events in which the Casein Kinase 1 (CK1) family plays a central, yet paradoxical, role. The long-standing model assumes a functional dichotomy: CK1α acts as a tumor suppressor by initiating β-catenin degradation, while CK1δ and CK1ɛ act as oncoproteins by promoting signalosome activity. However, their functionality is more complex, as for each of these isoforms positive as well as negative regulatory effects were described (see (Cruciat, 2014) for comprehensive review). Our study was designed to resolve the molecular basis behind the distinct roles of CK1α and CK1δ/ε and explore the potential for functional redundancy among these highly homologous kinases.

We confirm the established dogma of CK1α as a net negative regulator and CK1δ/ε as the net positive regulators in our CRISPR/Cas9 knockout (KO) models. Regarding the remaining CK1 family members, we previously showed that CK1α-like is not redundant with CK1α (Gybeľ et al., 2024). Probably the most surprising was our model of CK1γ1/2/3 triple KO, which lacked significant functional phenotype. These results are in collision with previous studies (Agajanian et al., 2022; Davidson et al., 2005) which reported CK1γ as a positive regulator of signaling through the phosphorylation of LRP5/6. Despite the individual limitations of these studies (overexpression, incomplete silencing, dependency on autocrine Wnt), it is too early to conclude that CK1γ does not play a significant role in the Wnt/β-catenin signaling. It is possible that in the absence of CK1γ paralogs, other CK1 kinases step in and replace CK1γ, for example in the phosphorylation of LRP5/6, which were previously implicated in this process (Dale, 2006). In fact, our study provides a proof of principle for this phenomenon in the destruction complex where CK1δ/ε replace CK1α in its absence. Interestingly, the recent study on CK1γ interactome identified also the components of Wnt/PCP signaling pathway (e.g. PRICKLE1, CELSR2, VANGL1/2) (Agajanian et al., 2022) as well as new PCP components RAI14 and EPHA2 (Gomoryova et al., 2025). This is in alignment with *Drosophila* CK1γ homolog gilgamesh (gish) and its role in Wnt/PCP, polarized vesicle trafficking (Gault et al., 2012). We also observed (data not shown) 50% decrease in the signaling capacity of CK1γ1/2/3 triple KO cells in Dual luciferase assay with ROR2/LRP6-chimeric receptor (Harada et al., 2017). The CK1γ subfamily will require further attention to elucidate its involvement in both Wnt/β-catenin and Wnt/PCP branches, position strikingly similar to CK1δ/ε.

A central question of our study has been how highly similar kinases can physiologically mediate opposing pathway outcomes. The overall sequence homology between CK1 isoforms does not intuitively explain their different roles; for instance, the functionally distinct CK1α and CK1α-like are more homologous than the functionally redundant CK1δ and CK1ɛ. Our *in vitro* kinase assays demonstrated, in line with earlier studies (Piao et al., 2011; Sakanaka, 2002), that CK1α and CK1ɛ do not exhibit significant intrinsic selectivity for canonical Wnt substrates like β-catenin or DVL3. This finding strongly suggests that their distinct functions *in vivo* are not determined by inherent enzymatic preference but rather by their integration into different protein complexes, guided by specific protein-protein interactions.

By employing a systematic chimera-based approach, we successfully mapped these functional determinants to distinct regions of the kinases. We identified the N-terminal lobe of CK1α as the critical element for its role within the β-catenin destruction complex. Swapping this region onto CK1ɛ conferred the ability to negatively regulate β-catenin, while removing it from CK1α abrogated this function. Conversely, we pinpointed the C-terminal tail of CK1ɛ as essential for its positive regulatory role at the DVL-dependent signalosome. Although this C-terminal domain is known to have an autoinhibitory function (Cegielska et al., 1998; Gietzen and Virshup, 1999; Rivers et al., 1998), our data clearly show it is also indispensable for productive signaling with DVL3, likely by mediating crucial protein interactions that scaffold the signalosome. Our results thus elaborate on previous *in vitro* study (Dahlberg et al., 2009) on CK1ε, indicating that two residues, N275 and R279, are crucial for the binding of Dishevelled (Dsh). Notably, in CK1α, the corresponding residues (I283 and T287) differ chemically, preventing such strong binding. These residues lie at the boundary of the region that differs between the long and short C- terminal swaps (see **Fig.** 3)–with one of these residues (CK1ε: R279, CK1α: T287) being replaced in the long C-terminal swap. Importantly, these long and short C-terminal swaps show a different effect on the signalosome function, providing independent support for the possible importance of R279 in DVL binding.

Furthermore, we suggest a mechanistic link for the N-terminal-driven specificity of CK1α by identifying SACK1G (also known as FAM83G and PAWS1) as a key interaction partner. Our co-immunoprecipitation experiments showed that the ability of our chimeric mutants to function in the degradasome assay perfectly correlated with their ability to bind SACK1G, an interaction dictated by the CK1α N-terminus. This aligns with recent studies emphasizing the importance of the CK1α-SACK1G axis in orchestrating the destruction complex (Bozatzi et al., 2018; Glennie et al., 2024) and provides a possible molecular basis for how CK1α is selectively recruited to its site of action.

Additionally, the identified CK1α-SACK1G binding mode correlates with observations from our previous study on CK1α-splice variants (Gybeľ et al., 2024). The L-segment of CK1α-v1 and v12 weakens the interaction with SACK1G (**Fig.** S8C/D, this study), which correlates with the weaker functionality of these variants in the degradasome assay (**Fig.** 4B in (Gybeľ et al., 2024)). Interestingly, CK1α-v2 as well as CK1α-like both bind SACK1G and phosphorylate β-catenin *in vitro*; however, only CK1α-v2 acts as a negative regulator, implying additional requirements, possibly the previously described activity towards Axin1 (Fulcher et al., 2018; Gybeľ et al., 2024).

Perhaps our most striking discovery is the functional plasticity of CK1δ and CK1ɛ. While they are established positive regulators of Wnt/β-catenin pathway, we found that in the absence of CK1α, they undergo a functional switch. Our data from pharmacological inhibition, genetic ablation, and acute protein degradation consistently show that, in CK1α-deficient cells, CK1δ/ɛ can functionally substitute for CK1α within the destruction complex, thereby becoming negative regulators of Wnt signaling. This was confirmed mechanistically by co-immunoprecipitation, which showed that CK1δ and CK1ɛ physically associate with the scaffold protein Axin1 only in cells lacking CK1α.

This finding fundamentally redefines the concept of redundancy for these kinases. It is not a simple case of one isoform compensating by performing the same task. Instead, it is a context-dependent rewiring where CK1δ/ɛ are co-opted for a novel function—the priming phosphorylation of β-catenin—when the primary kinase, CK1α, is absent. This resolves a significant contradiction in the field and explains why the outcome of modulating CK1 activity can be so context-dependent. Under normal conditions, CK1δ/ɛ inhibition dampens Wnt signaling by disrupting the signalosome. However, in a CK1α-deficient background, the same inhibition paralyzes the destruction complex, leading to a paradoxical and robust activation of the Wnt/β-catenin pathway.

Our findings could have profound implications for the development of CK1-targeted therapies. This work provides a framework for designing next-generation, isoform-selective CK1 modulators. By identifying the N- and C-terminal regions as determinants of function, we have highlighted novel surfaces that could be targeted to disrupt specific protein-protein interactions (e.g., the CK1α-SACK1G interface), rather than inhibiting the highly conserved kinase domain. While this study was conducted in HEK293 cells, future work should validate this functional switching model in relevant cancer cell lines and *in vivo* models. Investigating the precise structural motifs within the N- and C-terminal regions that mediate these specific interactions will be essential for developing truly targeted therapeutics.

In conclusion, we not only confirm the established dogma in our CRISPR/Cas9 knockout models but also uncover the structural determinants of this functional specificity and reveal a novel, context-dependent functional plasticity that fundamentally reframes our understanding of CK1 redundancy. This clarified understanding is not only a significant biological insight but also a crucial step toward the safe and effective therapeutic targeting of the CK1 family in human disease.

## MATERIALS & METHODS

### Cell culture

The T-REx-293 cell line (Invitrogen, R71007), Flp-In T-REx-293 (Invitrogen, R78007), and all derived cell lines were maintained in high-glucose Dulbecco’s Modified Eagle Medium (DMEM, Gibco, 41966029) supplemented with 10 % (v/v) fetal bovine serum (FBS, Gibco, A5256701) and 1 % (v/v) penicillin/streptomycin (Biosera, XC-A4122/100), further referred to as complete DMEM. Cells were cultivated in the cell culture incubator at 37 °C under 95 % (v/v) humidity and 5 % (v/v) CO_2_. Mycoplasma absence was routinely verified via PCR using forward primer: TYCTACGGGAGGCAGCAG and reverse primer: CGRCTGCTGGCACATAGTT (Siegl et al., 2023). When needed, a monolayer of cells was rinsed with phosphate-buffered saline (PBS; 140 mM NaCl; 8 mM Na_2_HPO_4_.12H_2_O; 1 mM KH_2_PO_4_; distilled H_2_O) and afterwards 1x Trypsin/EDTA (Biosera, XC-T1717/100) was gently applied to the cells. After approximately 5 minutes of trypsinization in the incubator, 1x Trypsin/EDTA was neutralized by addition of complete DMEM, and the obtained suspension of cells was subsequently handled and cultured on appropriate types of plates (TPP Techno Plastic Products AG or Thermo Scientific Nunclon Delta Surface) according to experimental setup or routine maintenance.

### CRISPR/Cas9 knockout cell lines

The procedure was described previously in (Gybeľ et al., 2024). Briefly, the design of gRNA sequences was carried out using the CRISPick-CRISPRko online designing tool, accessible at https://portals.broadinstitute.org/gppx/crispick/public (Doench et al., 2016; Sanson et al., 2018). For each gRNA, two oligonucleotides were designed (Table 1), hereafter referred to as the top and bottom oligos.

**Table 1:**
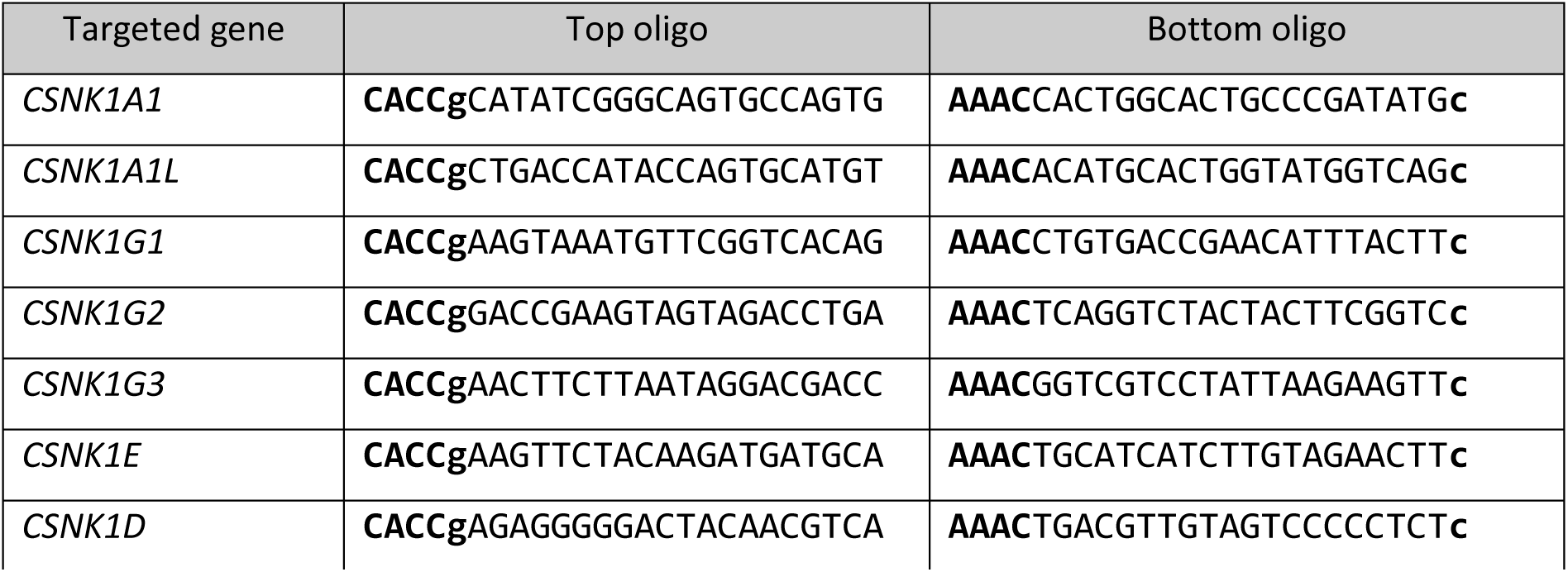
List of top and bottom oligos used for cloning of individual gRNAs.

The top oligo contains the exact gRNA sequence, while the bottom oligo is its reverse complement. Both oligos include additional nucleotides (in bold) to facilitate optimal annealing and cloning into one of the vector backbones. These express fluorescently tagged Cas9 nuclease and a single gRNA: pSpCas9(BB)-2A-GFP, addgene #48138 (Ran et al., 2013); pU6-(BbsI)_CBh-Cas9-T2A-mCherry, addgene #64324 (Chu et al., 2015); pU6-(BbsI)_CBh-Cas9-T2A-BFP, addgene #64323 (Chu et al., 2015). Cloning was performed according to the protocol described in Ran et al. 2013. The prepared vectors were validated via Sanger sequencing using a primer specific to the U6 promoter region: 5’-GAGGGCCTATTTCCCATGATTCC-3’.

Plasmids were transfected alone or in combination using Lipofectamine 2000 (Invitrogen, 11668027) according to the manufacturer’s instructions. The next day, cells underwent fluorescence-activated cell sorting (FACS) using a BD FACSAria Fusion system to establish monoclonal cell lines. Cells were expanded until they reached sufficient density. A fraction of each population was used for the isolation of genomic DNA (gDNA) with DirectPCR Lysis Reagent (Cell) (Viagen Biotech, 302-C) and 0.5 mg/ml Proteinase K (Thermo Scientific, EO0491) according to the manufacturer’s instructions. Isolated gDNA served as a template in a PCR reaction involving DreamTaq DNA Polymerase (Thermo Scientific, EP0702) and primers (Table 2) flanking the targeted locus.

**Table 2:**
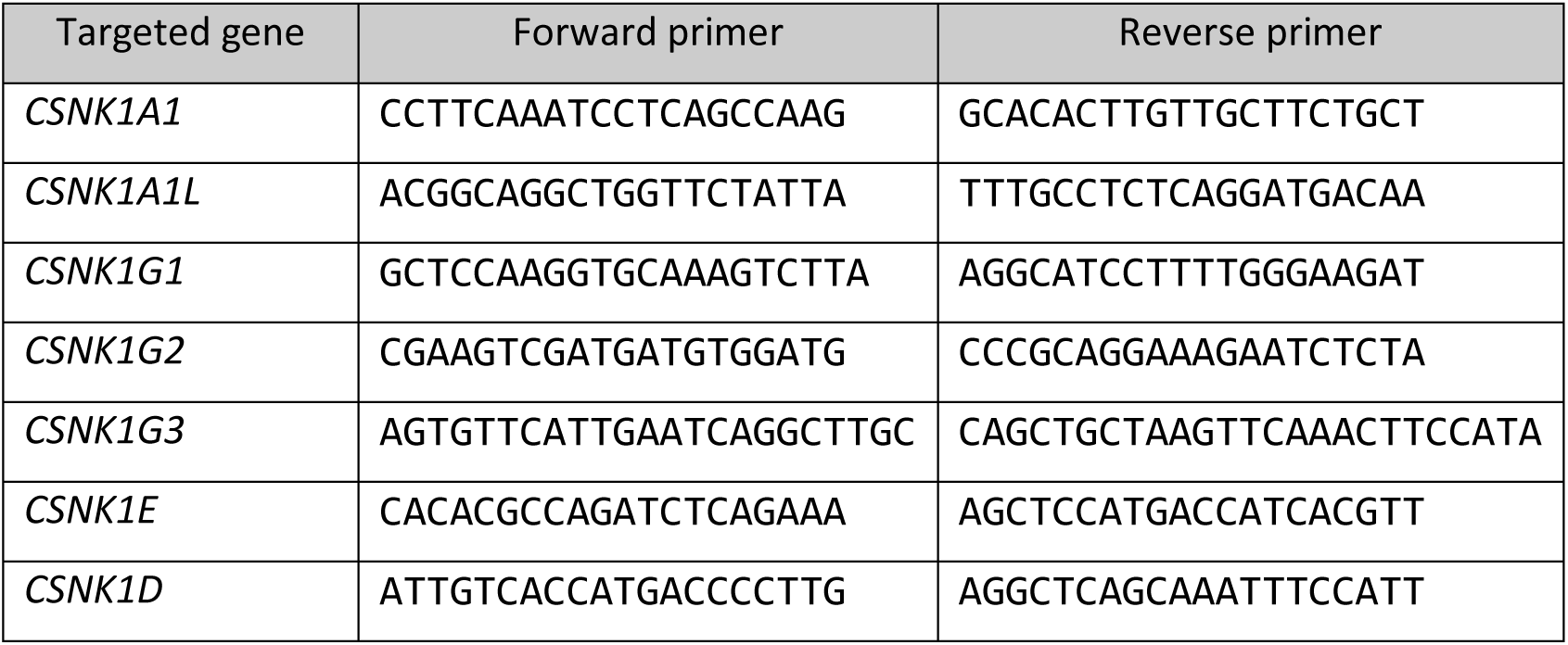
List of forward and reverse primer pairs for amplification of CRISPR/Cas9 targeted loci. Primers for *CSNK1G2* provided only a limited amount of PCR product, suitable only for NGS.

The obtained PCR product underwent simplified Restriction Fragment Length Polymorphism (RFLP) analysis to narrow down potential KO clones. Effective CRISPR/Cas9 modification with the selected gRNA should cause indels potentially disrupting the recognition site of the restriction enzyme (**Fig.**S1A; **Fig.**S2A), resulting in an uncleaved PCR product. RFLP analysis was visualized on a 2 % (w/v) agarose gel (Serva, 11404) alongside ZipRuler Express DNA Ladder Set (Thermo Scientific, SM1373). As some modifications may be present without affecting the restriction enzyme recognition site, clones were further or simultaneously submitted for WB analysis. Final confirmation was provided by Next Generation Sequencing (NGS) of the PCR product on the MiSeq Illumina platform following the previously described protocol (Malcikova et al., 2015). Analysis was carried out using the Integrative Genomics Viewer software – IGV (Robinson et al., 2011).

### Co-immunoprecipitation

For Flag-CK1–SACK1 co-immunoprecipitations, 10 cm plates of transfected T-REx-293 cells were washed once with room-temperature PBS and dry-frozen on dry ice before transferring to -80 °C for at least one hour. The following work was performed at 4 °C. Plates were placed on ice, and 1 ml of lysis buffer per plate (50 mM Tris-HCl at pH 7.4; 270 mM sucrose; 150 mM NaCl; 1 mM EDTA at pH 8.0; 1 mM EGTA at pH 8.0; 1 mM sodium orthovanadate; 10 mM sodium β-glycerophosphate; 50 mM sodium fluoride; 5 mM sodium pyrophosphate; 1 % (v/v) Nonidet P40 substitute) supplemented with 1x Protease inhibitor cocktail (PIC) (Roche, 11873580001) was added. Lysis was carried out for 15 minutes. Crude cell lysates were collected into tubes and sonicated on ice (7 s at 20 % amplitude) before clearance at 17,000 x g for 30 minutes. Protein concentration was determined using Bradford-based Protein Assay Dye Reagent (Bio-Rad, #5000006). Each sample was adjusted to 2 mg per 500 μl with lysis buffer. From this volume, 50 μl was taken as total cell lysate (TCL) input and mixed with 20 μl of NuPAGE 4x LDS sample buffer (Invitrogen, NP0007) containing 10 % (v/v) β-mercaptoethanol. The remaining volume was incubated with 12.5 μl of solid Anti-FLAG M2 conjugated beads (Sigma-Aldrich, A2220) for 2 h at 4 °C on a rotating wheel. After incubation, beads were pelleted (0.1 x g for 1 minute), and the supernatant (flow-through) was collected as a depletion control during optimization steps. Beads were washed three times with lysis buffer without PIC. The washed solid beads were resuspended in 25 μl of NuPAGE 2x LDS sample buffer containing 5 % (v/v) β-mercaptoethanol and stored at -20 °C. Samples were boiled for 5 minutes at 95 °C and either centrifuged at 16,000 x g for 1 minute to pellet beads or filtered using Spin-X (Costar, 8161) to remove beads before loading onto SDS-PAGE gel.

For endogenous co-immunoprecipitations, 10 cm plates of T-REx-293 cells were washed twice with room-temperature PBS and dry-frozen on plates at -80 °C for at least one hour. The following work was performed at 4 °C. Plates were placed on ice, and 1 ml of lysis buffer (50 mM Tris-HCl at pH 7.5; 150 mM NaCl; 1 mM EDTA; 0.5 % (v/v) Nonidet P40) supplemented with 1x Protease inhibitor cocktail (PIC) (Roche, 11836145001), 1mM dithiothreitol (DTT; Sigma-Aldrich, 43819), Phosphatase inhibitor cocktail I (1:100; Sigma-Aldrich, P2850) and Phosphatase inhibitor cocktail set II (1:100; Merck, 524625) was added. Lysis was carried out for 30 minutes. Crude cell lysates were collected into tubes and cleared at 14,000 x g for 20 minutes. Protein concentration was determined using the DC protein assay kit II (Bio-Rad, # 5000112). Each sample was adjusted to 2 mg per 700 μl with lysis buffer. From this volume, 60 μl was taken as a total cell lysate (TCL) input and mixed with 15 μl of 5x Laemmli buffer. The remaining lysate was incubated overnight at 4 °C with 1 μg of antibody (CK1ε – BD Biosciences 610445; CK1α – abcam ab206652; CK1δ – Santa Cruz Biotechnology sc-55553; Axin1 – Cell Signaling Technology #3323) or the corresponding isotype control (Mouse IgG1 – Cell Signaling Technology #5415; Rabbit IgG – Cell Signaling Technology #3900; Normal mouse IgG – Santa Cruz Biotechnology sc-2025). Protein G sepharose beads (GE Healthcare, GE17- 0618-05) were washed twice with lysis buffer and blocked overnight with 1 % (w/v) BSA in PBS. The following day, beads were washed three times with lysis buffer and once with lysis buffer containing supplements. Lysates with antibodies were incubated with 25 μl of solid beads for 7 hours at 4 °C on a rotator. After incubation, beads were pelleted (0.1 x g for 1 minute), and the supernatant (flow-through) was collected as a depletion control during optimization steps. Beads were washed four times with lysis buffer without supplements. Washed solid beads were resuspended in 60 μl of 2x Laemmli buffer and stored at -20 °C. Samples were boiled for 5 min at 95 °C and centrifuged to pellet beads before loading onto SDS-PAGE gel.

### Plasmids, Cloning, and Mutagenesis

Mammalian expression vectors of human CK1α and its variants (in detail described in (Gybeľ et al., 2024)), CK1α-like, CK1δ, and CK1ε were constructed using Gateway cloning. Original entry clones (Johannessen et al., 2010), pDONR223-CSNK1A1 (addgene, #23355), pDONR223-CSNK1A1L (addgene, #23784), pDONR223-CSNK1E (addgene, #23797), and pDONR223-CSNK1D (addgene, #23796) were adjusted using QuikChange II XL site-directed mutagenesis kit (Agilent, 200522) to repair the STOP codon for N-terminal fusion. Vector pDONR223-CSNK1D contained a silent mutation in the ORF of CSNK1D c.1077C>T, which remained uncorrected.

Construction of expression clones was carried out using prepared entry clones and the destination vector pDEST-pcDNA3.1 N-term 3xFlag in the reaction with the Gateway LR Clonase II Enzyme mix (Invitrogen, 11791020).

Further adjustments in terms of deletions or point mutations were carried out using the QuikChange II XL site-directed mutagenesis kit (Agilent, 200522). More extensive rearrangements (as in the case of CK1 chimeras) were performed using In-Fusion HD EcoDry Cloning Plus (Takara Bio USA, Inc.; 638913) and gene fragments (GENEWIZ, FragmentGENE).

Plasmids used in TurboID experiments are specified in the “TurboID” section. Plasmids used in the production of KO lines are specified in the “CRISPR/Cas9 knockout cell lines” section. Plasmids used in TOPFlash experiments are specified in the “Dual luciferase assay” section.

The sequences of all constructs in this study were verified by Sanger sequencing (Eurofins, LIGHTrun) and/or whole plasmid sequencing (Eurofins, Oxford Nanopore Technology). Sequences of cloning and mutagenesis primers, as well as gene fragments, are available upon request.

### SDS-PAGE and Western Blotting

Protein samples were, if not stated otherwise, prepared by direct lysis of cells with 2x Laemmli buffer (4 % (w/v) SDS; 20 % (v/v) glycerol; 0.125 M Tris-HCl at pH 6.8; 10 % (v/v) β-mercaptoethanol; 0.004 % (w/v) bromphenol blue), sonication and subsequent boiling of samples for 5 minutes at 95 °C.

#### SDS-PAGE and Western blotting were performed using established protocols

SDS-PAGE protein separation was carried out in several formats. Different densities of Tris-Glycine polyacrylamide gels were used in the Bio-Rad and ATTO systems. Additionally, NuPAGE 4-12 % Bis-Tris Midi Gels (Invitrogen, WG1403BX10) were used in the Invitrogen system. Electrophoresis was performed in the 1x Running buffer (190 mM glycine; 25 mM Tris; 0.1 % (w/v) SDS) for Tris-Glycine gels or 1x MOPS buffer for Bis-Tris gels at 80-150 V until the desired resolution was achieved. PageRuler Plus Prestained Protein Ladder (Thermo Scientific, 26620) was used as a size standard.

Western blot transfer to the methanol-activated polyvinylidene difluoride membranes (Immobilon-P PVDF, Millipore, IPVH00010) was carried out in 1x Transfer buffer (190 mM glycine; 25 mM Tris; 20 % (v/v) ethanol) at 105 V for 70 minutes. Alternatively, transferred membranes were stained with Ponceau S staining solution (Thermo Scientific, A40000278) to visualize successful transfer and the absence of bubbles. Membranes were blocked in 5 % (w/v) skimmed milk in TBST buffer (100 mM NaCl; 10 mM Tris-HCl at pH 7.6; 0.08 % (v/v) Tween20) for 1 hour at room temperature. Primary antibodies (Table 3) were diluted in 5 % (w/v) skimmed milk in TBST buffer and incubated with membranes overnight at 4 °C. The following day, membranes were washed three times for 15 minutes in TBST buffer and incubated with peroxidase-conjugated secondary antibodies (Anti-Rabbit IgG, Sigma-Aldrich A6667; Anti-Mouse IgG, Sigma-Aldrich A6782) diluted 1:5000 in 5 % (w/v) skimmed milk in TBST buffer for 1 hour at room temperature. Membranes were washed three times for 15 minutes in TBST buffer and covered with chemiluminescent HRP substrate (Millipore, WBKLS0500 or Bio-Rad, 170-5061) according to the manufacturer’s instructions. Emitted light was captured in a camera chamber, Fusion SL Vilber Lourmat or ChemiDoc MP Imaging System, using provided software and lossless TIFF (Tagged Image File Format) file format.

**Table 3:** List of primary antibodies used for Western blotting analysis.

| Primary Antibody anti- | Producer company | Catalog number | Working dilution | Species |
| --- | --- | --- | --- | --- |
| α-tubulin | Sigma-Aldrich | T6199 | 1:2000 | Mouse |
| β-actin | Cell Signaling | cs-4970 | 1:2000 | Rabbit |
| AXIN1 | Cell Signaling | cs-2087 | 1:1000 | Rabbit |
| Active β-catenin (depS37/T41) | MERCK Millipore | 05-665 | 1:500 | Mouse |
| Inactive β-catenin (pS45/T41) | Cell Signaling | cs-9565 | 1:1000 | Rabbit |
| total β-catenin | BD Biosciences | 610153 | 1:2000 | Mouse |
| Phospho-β-catenin (pS45) | Cell Signaling | cs-9564 | 1:1000 | Rabbit |
| CK1α | abcam | ab206652 | 1:1000 | Rabbit |
| CK1δ | Santa Cruz | sc-55553 | 1:200 | Mouse |
| CK1ε | BD Biosciences | 610445 | 1:2000 | Mouse |
| CK1γ1 | abcam | ab234727 | 1:1000 | Rabbit |
| CK1γ2 | Santa Cruz | sc-390970 | 1:500 | Mouse |
| CK1γ3 | Novus Biologicals | NBP1-57573 | 1:100 | Rabbit |
| CK1γ3 | Sigma-Aldrich | HPA027010 | 1:100 | Rabbit |
| CK1γ3 | abcam | ab37950 | 1:100 | Rabbit |
| DVL2 | Cell Signaling | cs-3216 | 1:1000 | Rabbit |
| DVL3 | Cell Signaling | cs-3218 | 1:1000 | Rabbit |
| DVL3 | Santa Cruz | sc-8027 | 1:1000 | Mouse |
| SACK1B (FAM83B) | MRC PPU Reagents and Services | DU28279 (SA270) | 1 µg/ml | Sheep |
| SACK1G (FAM83G) | abcam | ab121750 | 1:1000 | Rabbit |
| SACK1H (FAM83H) | MRC PPU Reagents and Services | DU28403 (SA273) | 1 µg/ml | Sheep |
| Flag | Sigma-Aldrich | F1804 | 1:2000 | Mouse |
| Flag | Sigma-Aldrich | F7425 | 1:2000 | Rabbit |
| Flag-HRP | Sigma-Aldrich | A8592 | 1:10 000 | (Mouse) |
| GAPDH | Cell Signaling | cs-5174 | 1:2000 | Rabbit |
| GAPDH-HRP | Proteintech | HRP-60004 | 1:10 000 | (Mouse) |
| GSK3α/β | Cell Signaling | cs-5676 | 1:1000 | Rabbit |
| HA | abcam | ab9110 | 1:2000 | Rabbit |
| HA | Biolegend | 901514 | 1:2000 | Mouse |
| LRP6 pS1490 | Cell Signaling | cs-2568 | 1:500 | Rabbit |
| LRP6 total | Cell Signaling | cs-3395 | 1:1000 | Rabbit |
| Phospho-VANGL2 (pT78/S79/S82) | ABclonal | AP1206 | 1:1000 | Rabbit |
| Vinculin | Cell Signaling | cs-4650 | 1:1000 | Rabbit |

Re-probing the membrane for an additional primary antibody of a different species, the membrane was first treated with a quenching buffer composed of 10 % (v/v) acetic acid for 30 minutes at 37°C (Han et al., 2020). In the case of re-probing for an additional primary antibody of the same species, the previous antibody was stripped using a guanidine hydrochloride (GnHCl)-based stripping buffer for 30 minutes at room temperature (Yeung and Stanley, 2009).

ImageJ software (Schneider et al., 2012) was used for the post-processing of images in terms of their cropping and linear adjustments of brightness and contrast.

### Transient transfection

In case of TOPFlash: degradasome assay, transient transfections were performed using polyethylenimine (PEI, Polysciences, 23966) as previously described in (Gybeľ et al., 2024). Briefly, cells were plated on a 24-well plate a day in advance to be on the day of transfection at 60-70 % confluency.

The total volume of serum/antibiotics-free DMEM used was 50 μl. Half of this volume (25 μl) was incubated with 1.2 μl of PEI (1 µg/ml; pH 7.4) for 30 minutes, while the other half (25 μl) was mixed with the desired DNA for transfection. The total amount of DNA was 0.4 μg, equilibrated by pcDNA3 empty backbone (Invitrogen) when necessary. The resulting DNA:PEI ratio was 1:3 (w/v). Once prepared, both mixtures were combined and incubated for 30 minutes. Mixture was added to the cells and incubated for 5 hours. Afterwards, the medium was replaced with fresh complete DMEM, alternatively with treatment.

Transient transfections for the TOPFlash: signalosome assay were carried out similarly; however, PEI was replaced with jetOPTIMUS (Polyplus, 101000025) at a DNA:jetOPTIMUS ratio of 1:1 (w/v).

Transient transfections for Flag-CK1–SACK1 co-immunoprecipitations were performed using PEI (Sigma-Aldrich, 919012) in 10 cm plates. The total transfected DNA amount was 10 μg (5 μg of expression construct and 5 μg of empty pcDNA3 backbone), with a DNA:PEI ratio of 1:2 (w/v) in a total transfection mixture volume of 1000 μl. Transfection was conducted overnight.

### Dual luciferase assay

The dual luciferase assay for measuring the canonical Wnt signaling pathway is commonly referred to as the TOPFlash/Renilla assay or simply TOPFlash. This method employs Firefly and Renilla luciferases to comparatively quantify the expression of TCF/LEF-responsive genes, which are regulated by the activation of the Wnt/β-catenin signaling pathway (Korinek et al., 1997).

The assay was described in more detail previously in Gybel et al., 2024. Briefly, cells were co-transfected (see Transient transfection) with two main vectors: 100 ng of Super 8x TOPFlash (Veeman et al., 2003) and 100 ng of pRL-TK vector (Promega) per 1 well of a 24-well plate. Additional vectors depended on the experimental setup. For the TOPFlash: degradasome assay, additional vectors were 50ng of CK1 isoforms or their mutants. In the case of TOPFlash: signalosome assay, additional vectors were 10 ng of pCMV-HA-hDvl3 and/or 150 ng of CK1 isoforms or their mutants. Cells were transfected for 5 hours and subsequently treated overnight (16 hours) according to experimental setup with 0.5 µM LGK974 (Stem RD, L974-010), WNT3A conditioned medium (WNT3A CM), 80ng/ml of WNT3A recombinant protein (WNT3A RP) (R&D Systems, 1324-WN) or 25ng/ml of R-spondin-1 (RSPO1 RP) (PeproTech, 120-38) or their combinations. Control treatments were control conditioned medium and 0.1 % (w/v) bovine serum albumin (BSA, Serva, 11930.03) in PBS, which was used as a solvent of recombinant proteins.

Dual luciferase assay was performed with the Dual-Luciferase Reporter Assay System (Promega, E1960), according to the manufacturer’s instructions. Briefly, the next day after the transfection, all liquid was aspirated, and cells were dry-frozen at -80 °C for at least 1 hour. Once plates reached room temperature, cells were lysed with 50 μl/well of passive lysis buffer for 20 minutes at room temperature while rocking the plate. Then, 20 μl/sample of lysate was moved to flat bottom strips (Thermo Scientific, 7566) and 25 μl/sample of Luciferase Assay Reagent II (LAR II) was added. Strips were inserted into the Hidex Bioscan Plate Chameleon Luminometer, and luminescence produced by Firefly luciferase (TOPFlash) was measured. In the next step, 25 μl/sample of Stop&Go reagent was added, and luminescence produced by Renilla luciferase was measured. Obtained raw luminescence units for the Firefly luciferase (TOPFlash) were divided/normalized by raw luminescence units for Renilla luciferase, providing relative luminescence units (RLU). Each RLU value of a given replicate was further normalized to the mean value of the whole replicate. Eventually, each value across replicates was normalized by the mean value of controls (T-REx-293 WT Ctrl/ pcDNA / -WNT3A, -RSPO1) across replicates. Data normalized in this way were plotted and represent fold changes in the relative activity of the Wnt/β-catenin signaling pathway. Graphs and statistical evaluation were carried out in GraphPad Prism 8 software using Two-Way or Three-Way ANOVA with Tukey’s multiple comparisons test.

### Treatments of cells

In this study, several types of treatments were used. To inhibit the autocrine secretion of WNT ligands, cells were treated with Porcupine inhibitor LGK974 (Stem RD, #L974-010) at a concentration of 0.5 µM for 16 hours. To stimulate Wnt/β-catenin signaling cascade, mouse recombinant WNT3A (R&D Systems, #1324-WN) was used at a concentration of 80 ng/ml for 16 hours. Alternatively, cells were treated with conditioned medium (CM), in 1:1 ratio with fresh complete DMEM, prepared culturing L cells (CRL-2648, ATCC) for CTRL CM or L Wnt-3A cells (CRL-2647) for WNT3A CM according to the manufacturers instructions. To potentiate the stimulation, recombinant protein R-spondin-1 (RSPO1, PeproTech, # 120-38) was used at a concentration of 25 ng/ml for 16 hours. Control stimulations were carried out with 0.1% bovine serum albumin (BSA, Serva, #11930.03) in PBS, which was used as a solvent of recombinant proteins.

The following inhibitors and degraders were used at the concentrations stated in the figures: PF-670462 (DC2086, DC Chemicals), CHIR-99021 (HY-10182, MedChem Express), XAV-939 (HY-15147, MedChem Express), MU1742 (a generous gift from the group of Kamil Paruch), DEG-77 (a generous gift from the group of Kamil Paruch; see (Park et al., 2023)), SJ3149 (HY-160444, MedChem Express), AH078 (a generous gift from the group of Stefan Knapp; see (Haag et al., 2025)).

### Protein expression and purification

Purification of hCK1α full-length (FL; 1-337 aa; CK1α-v2) (Gybeľ et al., 2024), hDVL3 (335-396 aa) (Micka *et al*., 2025; manuscript) and hCK1ε full-length (FL; 1-416 aa) (Bologna *et al*., 2025; manuscript) were described elsewhere.

Human β-catenin (1-145 aa) and VANGL2 (1-100 aa) were expressed in *E. coli* BL21(DE3) bacterial cells from pETM11 backbones containing N-terminal His- and Z-tags. Cells were grown at 37°C in M9 mineral medium until reaching an OD_600_ ∼ 0.7. Expression was induced by 0.25 mM IPTG, followed by 3-4 hours of cultivation at 37°C. Cells were then harvested, resuspended in lysis buffer (25 mM Tris-HCl, 500 mM NaCl, 300 mM sucrose, 10 % glycerol, 10 mM imidazole, 1 % Triton X-100, pH 8.0), and stored at -20°C. Defrosted cells were supplemented with protease inhibitor (AppliChem, PMSF), sonicated (amplitude 22) at 4 °C for 20 min (5 s ON / 10 s OFF), and lysate was clarified by centrifugation at 27,000 × g for 1 h at 4 °C.

In the case of β-catenin, immobilized metal affinity chromatography (IMAC) was performed via His₆-tag with a Ni-NTA column (Cytiva, HiTrap IMAC HP 5 ml). Filtered (0.45 μm filter) supernatant was loaded onto the sorbent. Unbound contaminants were washed out using binding buffer (25 mM Tris-HCl at pH 8.0, 500 mM NaCl, 10 mM imidazole, 5 % glycerol), and the protein of interest was afterwards eluted in a gradient of the binding buffer supplemented with 500 mM imidazole. The eluant was supplemented with 1mg/ml TEV protease (produced in-house), 2 mM β-mercaptoethanol, and 1 mM EDTA for 1 h at 25 °C to cleave the tag, and dialyzed to the binding buffer for 16 h at 4 °C to remove the imidazole. The next day, the sample was boiled, snap-cooled, and centrifuged at 4000 × g for 20 min at 25 °C before loading into the second IMAC for tag removal. The flow-through containing the peptide was concentrated by centrifugal ultrafiltration (Vivaspin 20, 5000 MWCO PES, Sartorius) and subjected to size-exclusion chromatography (SEC; Superdex 75 10/300 GL, Cytiva) in NMR buffer (50 mM phosphate buffer at pH 6.5, 50 mM KCl). Aliquots were snap-frozen in liquid nitrogen and stored at -80 °C.

Purification of VANGL2 was performed similarly to β-catenin, except for two steps. Dialysis was performed to a low salt buffer (25 mM Tris-HCl, 50 mM NaCl, 1 mM EDTA, 5 % glycerol, pH 8.0). Ion exchange chromatography (IEX) was used instead of the second IMAC. Supernatant was loaded on the column (HiTrap Q HP 5 ml, Cytiva) in low salt buffer and eluted in the same buffer supplemented with 1M NaCl.

### *In vitro* kinase assay and MALDI

*In vitro* kinase assays with abovementioned kinases (CK1α FL and CK1ε FL) and substrates (β-catenin (1- 145 aa), hDVL3 (335-396 aa) and VANGL2 (1-100 aa)) were carried out in the buffer containing 50 mM phosphate buffer at pH 6.5, 50 mM KCl, 10 mM MgCl_2_ and 2 mM ATP. The molar ratio was set to 1:50, 1 µM kinase and 50 µM substrate. Reactions with just substrates (without kinases) served as controls. Reaction time was 24 hours at room temperature, and reactions were stopped by boiling at 95°C for 6 minutes. Reactions were centrifuged at 20,000 x g for 6 minutes and submitted for Matrix-assisted laser desorption/ionization-time of flight mass spectrometry (MALDI-TOF MS).

MALDI-TOF mass spectra of the proteins were recorded on an Ultraflextreme instrument (Bruker Daltonics, Bremen, Germany) operated in the linear positive ion detection mode under FlexControl 3.4 software (Bruker Daltonics). Mass spectra were processed with FlexAnalysis 3.4 software (Bruker Daltonics). Ferulic acid (12.5 mg.mL-1 in a mixture of water:acetonitrile:formic acid 50:33:17, v/v) was used as MALDI matrix. The samples in a volume of 0.6 µl were mixed with 2.4 µl of the matrix solution, and 0.6 µl of this mixture were deposited onto a stainless steel MALDI target. External calibration of the mass spectra was performed by utilizing signals of the ionization forms of chicken lysozyme.

### TurboID

Starting vector to produce TurboID lines was prepared re-cloning the sequence of BirA* in HA-BirA*- pDEST-N-pcDNA5/FRT/TO (addgene, #118375) (Viita et al., 2019) with TurboID BirA* from V5-TurboID-NES_pCDNA3 (addgene, #107169) (Branon et al., 2018) using In-Fusion HD EcoDry Cloning Plus (Takara Bio USA, Inc.; 638913). Prepared vector HA-TurboID-BirA*-pDEST-N-pcDNA5/FRT/TO was used in Gateway cloning using Gateway LR Clonase II Enzyme mix (Invitrogen, 11791020) together with prepared CK1 entry clones described in “Cloning and mutagenesis” and DVL3 (DNASU, HsCD00043699). Flp-In T-REx-293 (Invitrogen, R78007) were transfected with prepared vectors and Flp-Recombinase (pOG44; Invitrogen, V600520) using jetOPTIMUS (Polyplus, 101000025). Upon selection with Hygromycin B (50ug/ml; Gibco, 10687010), cells were stained for viability and FACS sorted to establish monoclonal cell lines. Clones were subsequently tested for the inducible expression of the fusion protein of interest from the integrated construct and their ability to induce biotinylation.

The TurboID experiment was performed according to the previously published BioID protocol (Roux et al., 2018). TurboID cell lines were plated in 15 cm dishes with complete DMEM containing Tetracycline-free FBS (Biosera, FB-1001T/500). Cells were treated with 200 ng/ml tetracycline (Santa Cruz Biotechnology, sc-205858) and 1 μM LGK974 (Stem RD, L974-010) for 18 hours before lysis. Cells were treated either with control conditioned medium or Wnt3a conditioned medium for 1.5 hours before lysis. Biotin (58 μM; Santa Cruz Biotechnology, sc-204706A) was added for the last 30 minutes before lysis.

Medium was removed, and cells were washed twice with room temperature PBS. Monolayer or cells was lysed with 4.5 ml of lysis buffer (50 mM Tris-HCl at pH 7.4; 500 mM NaCl; 0.4 % (w/v) SDS; 2 % (v/v) Triton X-100; 1x Protease inhibitor cocktail (PIC) (Roche, 11836145001), 1mM dithiothreitol (DTT; Sigma-Aldrich, 43819)) for 15 minutes at 4°C. Lysates were collected into tubes and sonicated (amplitude 2) on ice for 10 seconds, inverted a few times, and sonicated again for 10 seconds. Lysis was allowed to continue on ice for an additional 15 minutes following lysate clearance at 16.500 x g for 15 minutes at 4°C. Supernatant was transferred to a clean tube, and 68 μl was taken as a total cell lysate (TCL) input and mixed with 20 μl of 5x Laemmli buffer. The remaining lysate was diluted 2x with 50 mM Tris-HCl at pH 7.4. Streptavidin Sepharose High Performance beads (30 μl of solid beads per sample; Cytiva, GE17-5113-01) were washed twice in 2x diluted lysis buffer and incubated with lysate overnight (23 hours) at 4°C on a rotator. The following day, beads were pelleted at 100 x g for 5 minutes at 4°C. Supernatant/flow-through (68 μl mixed with 20 μl of 5x Laemmli buffer) was collected as a depletion control, and the rest was discarded. The following steps were carried out at room temperature. Beads were washed as follows: twice with 1 ml of wash buffer 1 (2 % (w/v) SDS), once with 1 ml of wash buffer 2 (0,1 % (w/v) deoxycholic acid; 1 % (w/v) Triton X -100; 1 mM EDTA; 500 mM NaCl; 50 mM HEPES at pH 7.5), once with 1 ml of wash buffer 3 (0.5 % (w/v) deoxycholic acid; 0.5 % (w/v) NP-40; 1 mM EDTA; 250 mM LiCl; 10 mM Tris-HCl at pH 7.4) and twice with 1 ml of 50 mM Tris-HCl at pH 7.4. Washed beads were stored at -80°C till MS analysis. TurboID experiment was performed in four biological replicates.

### Software

Alignments in this article were created using Jalview 2.11.4.1 (Waterhouse et al., 2009). AlphaFold3 predictions (**Fig.** 3Aiii and **Fig.** S6Aii) were generated on the AlphaFold Server, accessible at https://alphafoldserver.com/ (Abramson et al., 2024), using the canonical sequences of CK1α (UniProt: P48729-1) and CK1ε (UniProt: P49674).

## DATA AVAILABILITY

The data supporting the findings of this article are available from the corresponding author.

## SUPPORTING INFORMATION

This article contains supporting information.

## ACKNOWLEDGEMENTS

We thank Mgr. Tomáš Loja, Ph.D. and MVDr. Jana Černá, Ph.D. for FACS; Mgr. Šárka Pavlová, Ph.D. and colleagues for NGS, and all members of OFIŽ, for their encouragement and support. We acknowledge the use of AI tools (ChatGPT and Perplexity) for language and grammar improvements of this manuscript.

## FUNDING AND ADDITIONAL INFORMATION

This study was supported by the Czech Science Foundation - Grant projects of excellence in basic research EXPRO-Molecular and functional analysis of casein kinase 1 biology (GX19-28347X); The project National Institute for Cancer Research (Programme EXCELES, ID Project No. LX22NPO5102) - Funded by the European Union – Next Generation EU; European Structural and Investment Funds, Operational Programme Research, Development and Education „Preclinical Progression of New Organic Compounds with Targeted Biological Activity” (Preclinprogres —CZ.02.1.01/0.0/0.0/16_025/0007381); the project CZ-OPENSCREEN: National Infrastructure for Chemical Biology (CZ-OPENSCREEN LM2023052); and Bader Philanthropies. CIISB, Instruct-CZ Centre of Instruct-ERIC EU consortium, funded by MEYS CR infrastructure project LM2023042 and European Regional Development Fund-Project „Innovation of Czech Infrastructure for Integrative Structural Biology“ (No. CZ.02.01.01/00/23_015/0008175), is gratefully acknowledged for the financial support of the measurements at the CEITEC Proteomics Core Facility. Computational resources were provided by the e-INFRA CZ project (ID:90254), supported by MEYS CR. T.G. is Brno Ph.D. Talent 2019-2022 scholarship holder, which was funded by the Brno city municipality. T.G. is an EMBO Scientific Exchange Grant holder, which enabled placement in the laboratory of G.P.S. and the establishment of a collaboration.

