## Supporting information for "Molecular mechanisms behind the functional (non)redundancy of CK1 paralogs in the Wnt pathway"

* Correspondence to:

Department of Experimental Biology

TABLE OF CONTENTS:

Figure S1

Figure S2

Figure S3

Figure S4

Figure S5

Figure S6

Figure S7

Figure S8

Figure S9

### Figure S1

A

| Region [aa] | | CK1α | CK1α-like | CK1γ1 | CK1γ2 | CK1γ3 | CK1δ | CK1ε |
|---|---|---|---|---|---|---|---|---|
| 1-337 | CK1α | 100.00 | | | | | | |
| 1-337 | CK1α-like | 90.50 | 100.00 | | | | | |
| 1-422 | CK1γ1 | 47.95 | 46.82 | 100.00 | | | | |
| 1-415 | CK1γ2 | 49.56 | 47.49 | 78.77 | 100.00 | | | |
| 1-447 | CK1γ3 | 49.85 | 48.25 | 76.50 | 76.94 | 100.00 | | |
| 1-415 | CK1δ | 68.39 | 65.05 | 47.80 | 48.69 | 45.45 | 100.00 | |
| 1-416 | CK1ε | 68.69 | 65.35 | 47.10 | 47.68 | 47.10 | 83.17 | 100.00 |

B

| Region [aa] | | CK1α | CK1α-like | CK1γ1 | CK1γ2 | CK1γ3 | CK1δ | CK1ε |
|---|---|---|---|---|---|---|---|---|
| 1-16 | CK1α | 100.00 | | | | | | |
| 1-16 | CK1α-like | 68.75 | 100.00 | | | | | |
| 1-43 | CK1γ1 | 31.25 | 31.25 | 100.00 | | | | |
| 1-45 | CK1γ2 | 31.25 | 25.00 | 46.67 | 100.00 | | | |
| 1-42 | CK1γ3 | 25.00 | 31.25 | 46.51 | 46.67 | 100.00 | | |
| 1-8 | CK1δ | 37.50 | 50.00 | 37.50 | 37.50 | 25.00 | 100.00 | |
| 1-8 | CK1ε | 50.00 | 62.50 | 37.50 | 25.00 | 37.50 | 87.50 | 100.00 |

C

| Region [aa] | | CK1α | CK1α-like | CK1γ1 | CK1γ2 | CK1γ3 | CK1δ | CK1ε |
|---|---|---|---|---|---|---|---|---|
| 17-285 | CK1α | 100.00 | | | | | | |
| 17-285 | CK1α-like | 92.57 | 100.00 | | | | | |
| 44-315 | CK1γ1 | 56.25 | 54.04 | 100.00 | | | | |
| 46-316 | CK1γ2 | 56.46 | 54.24 | 93.75 | 100.00 | | | |
| 43-313 | CK1γ3 | 57.56 | 54.98 | 94.49 | 94.10 | 100.00 | | |
| 9-277 | CK1δ | 78.44 | 73.98 | 59.19 | 58.67 | 60.52 | 100.00 | |
| 9-277 | CK1ε | 78.07 | 73.61 | 59.56 | 59.04 | 60.89 | 97.40 | 100.00 |

D

| Region [aa] | | CK1α | CK1α-like | CK1γ1 | CK1γ2 | CK1γ3 | CK1δ | CK1ε |
|---|---|---|---|---|---|---|---|---|
| 286-337 | CK1α | 100.00 | | | | | | |
| 286-337 | CK1α-like | 86.54 | 100.00 | | | | | |
| 316-422 | CK1γ1 | 11.11 | 17.24 | 100.00 | | | | |
| 317-415 | CK1γ2 | 19.23 | 19.23 | 54.21 | 100.00 | | | |
| 314-447 | CK1γ3 | 17.65 | 20.37 | 50.37 | 52.24 | 100.00 | | |
| 278-415 | CK1δ | 21.15 | 21.15 | 19.63 | 23.30 | 16.67 | 100.00 | |
| 278-416 | CK1ε | 23.08 | 23.08 | 18.80 | 20.18 | 20.15 | 55.40 | 100.00 |

% Percentage identity

lowest highest

**Figure S1: Homology between CK1 isoforms.**
Percentage identity was calculated based on pairwise alignments using the BLOSUM62 matrix from ClustalOWS alignments generated in Jalview 2.11.4.1.
**(A)** Overall homology: Calculation based on the alignment of full-length protein sequences.
**(B)** N-terminal homology: Calculation based on the alignment of N-terminal protein sequences within specified regions.
**(C)** Kinase domain homology: Calculation based on the alignment of protein sequences in the regions of kinase domains.
**(D)** C-terminal homology: Calculation based on the alignment of C-terminal protein sequences within specified regions.

### Figure S2

A

| Gene | Protein | gRNA + **PAM**, Restriction site | PCR | RE | RFLP |
|---|---|---|---|---|---|
| *CSNK1G1* | CK1γ1 | AAGTAAATGTTCGGTCACAG**AGG** | 682bp | Tsp45I | 420bp + 216bp + 46bp |
| *CSNK1G2* | CK1γ2 | GACCGAAGTAGTAGACCTGA**GGG** | No PCR product available for Bsu36I restriction. | | |
| *CSNK1G3* | CK1γ3 | AACTTCTTAATAGGACGACC**AGG** | 552bp | BstNI | 352bp + 200bp |

B

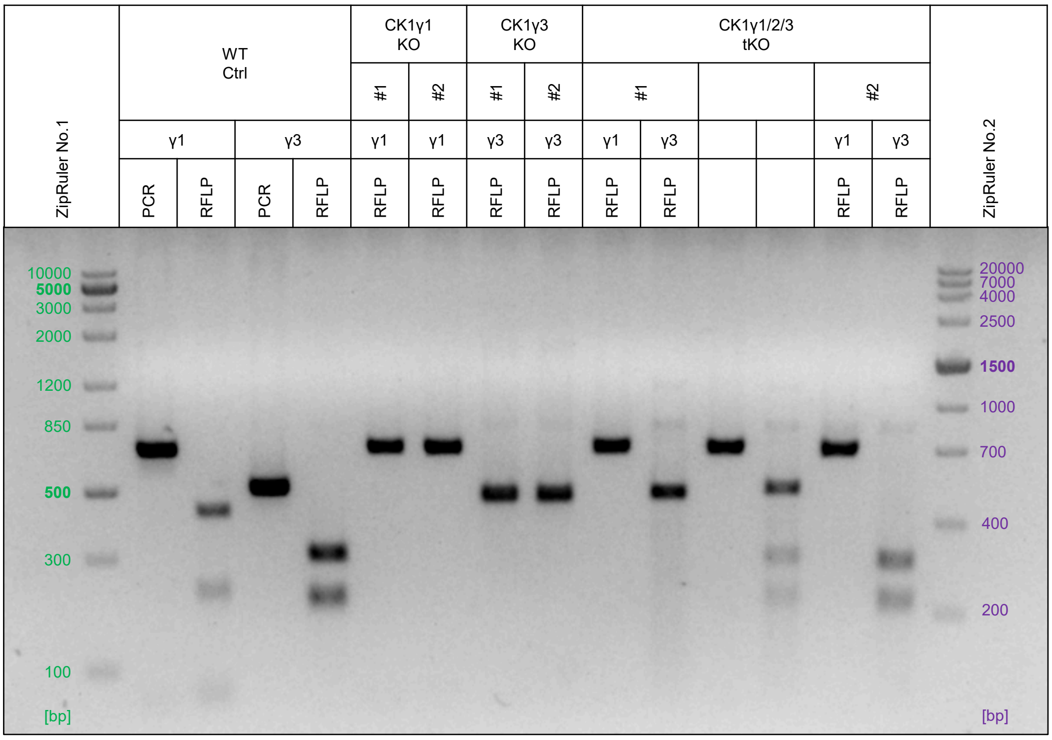


C

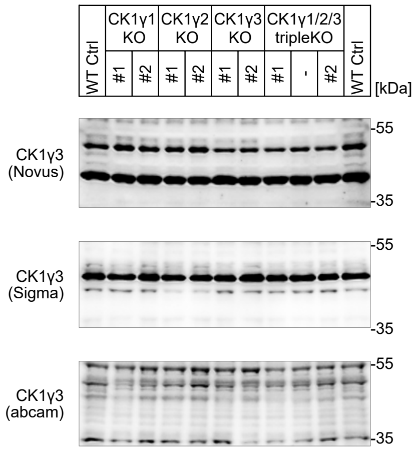


**Figure S2: Production and validation of CK1γ KO panel cell lines.**
**(A)** Sequences of gRNAs used to generate single and compound KO cell lines in the CK1γ KO panel (Fig.1Di). gRNAs were designed in accordance with PCR screening of KO clones using RFLP analysis. The PAM sequence (in bold) is not part of the gRNA sequence. Underlined sequences indicate recognition/cleavage sites of the specified restriction enzymes (RE). Expected PCR product sizes and corresponding restriction patterns are provided. Despite several optimizations, PCR product yield was insufficient for RFLP analysis of the targeted *CSNK1G2* locus; however, it was adequate for NGS sequencing (Fig.S2D).
**(B)** Agarose gel (2 %) showing RFLP analysis of KO clones.
**(C)** Extension of Fig.1Dii showing Western blot validation of the CK1γ KO panel using several CK1γ3 antibodies available at the time.

### Figure S2

D

| Next Generation Sequencing Results | | | | |
|---|---|---|---|---|
| *CSNK1G1* gene – hg19: chr15 (+) 64508788 - 64508820 | | | | |
| WT | | γ1 | cttcaAAGTAAATGTTCGGTCA-CAGAGGtcaaa | |
| *CSNK1G2* gene – hg19: chr19 (+) 1978603 - 1978635 | | | | |
| WT | | γ2 | ggcgtCCCTCAGGTCTACTACTTCGGTCcgtgc | |
| *CSNK1G3* gene – hg19: chr5 (+) 122911478 – 122911510 | | | | |
| WT | | γ3 | ctgagAACTTCTTAATAGGACGACCAGGaaacaaaaccca | |
| KO lines | | | | Type of modification |
| CK1γ1 KO | #1(F12) | γ1 | cttcaAAGTAAATGTTCGGTCAACAGAGGtcaaa | 1nt ins |
| | #2(C12) | γ1 | cttcaAAGTAAATGTTCGGTCAACAGAGGtcaaa<br>cttcaAAGTAAATGTTCGGTCA-CAGAGGtcaaa | 1nt ins<br>5nt del |
| CK1γ2 KO | #1(C5) | γ2 | ggcgtCCCTCAGGGTCTACTACTTCGGTCcgtgc | 1nt ins |
| | #2(G8) | γ2 | ggcgtCCCTCAGGGTCTACTACTTCGGTCcgtgc | 1nt ins |
| CK1γ3 KO | #1(26) | γ3 | ctgagAACTTCTTAATAGGACGACCAGGaaacaaaaccca | 4nt del |
| | #2(29) | γ3 | ctgagAACTTCTTAATAGGACGACCAGGaaacaaaaccca<br>ctgagAACTTCTTAATAGGACGACCAGGaaacaaaaccca | 2nt del<br>8nt del |
| CK1γ1/2/3 tripleKO | #1(7) | γ1 | cttcaAAGTAAATGTTCGGTCAACAGAGGtcaaa | 1nt ins |
| | | γ2 | ggcgtCCCTCAGGGTCTACTACTTCGGTCcgtgc<br>ggcgtCCCTCAGGTCTACTACTTCGGTCcgtgc | 1nt ins<br>11nt del |
| | | γ3 | ctgagAACTTCTTAATAGGACGACCAGGaaacaaaaccca | 15nt + 13nt del |
| | #2(15) | γ1 | cttcaAAGTAAATGTTCGGTCAACAGAGGtcaaa | 1nt ins |
| | | γ2 | ggcgtCCCTCAGGTCTACTACTTCGGTCcgtgc | 1nt del |
| | | γ3 | ctgagAACTTCTTAATAGGACGACCAGGaaacaaaaccca | 10nt del |

E

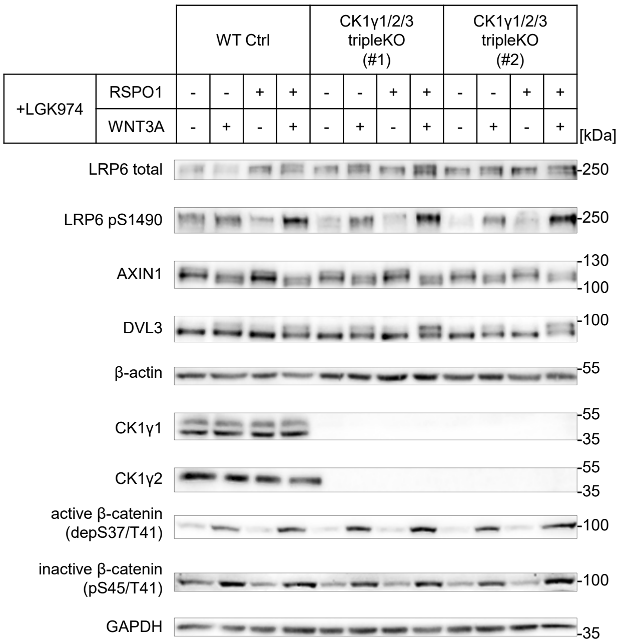


**Figure S2: Production and validation of CK1γ KO panel cell lines.**
**(D)** NGS results confirming the KO status of the presented cell lines. PAM sequences are displayed in blue, orange, and purple for the individual CK1γ isoforms, while gRNA sequences are displayed in black uppercase letters. Multiple sequences within one isoform represent different allele modifications. Dashes are included to improve sequence alignment. ins = insertion; del = deletion.
**(E)** Western blot analysis of CK1γ1/2/3 tripleKO clones #1 and #2 using protein lysates from the TOPFlash analysis (Fig.1Diii).

### Figure S3

A

| Gene | Protein | gRNA + **PAM**, Restriction site | PCR | RE | RFLP |
|---|---|---|---|---|---|
| *CSNK1E* | CK1ε | AAGTTCTACAAGATGATGCA**GGG** | 396bp | SfaNI | 215bp + 181bp |
| *CSNK1D* | CK1δ | AGAGGGGGACTACAACGTCA**TGG** | 369bp | DrdI | 228bp + 141bp |

B

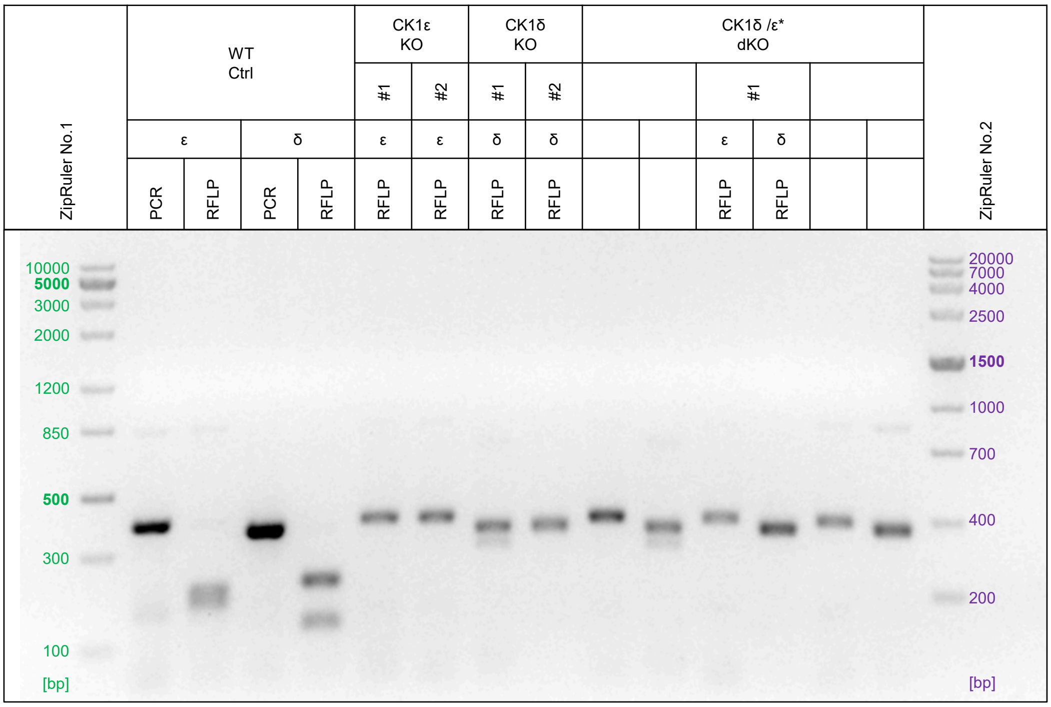


C

| Next Generation Sequencing Results | | | | |
|---|---|---|---|---|
| *CSNK1E* gene - hg19: chr22 (+) 38699143 - 38699175 | | | | |
| WT | | ε | accgccaCCCTGC--ATCA---TCTTGTAGAACTTgctct | |
| *CSNK1D* gene - hg19: chr17 (+) 80213395 - 80213427 | | | | |
| WT | | δ | gcagctccatca-CCATGA-CGTTGTAGTCCCCCTCTgcccc | |
| KO lines | | | | Type of modification |
| CK1ε KO | #1(ε4) | ε | accgccaCCCTGCA-ATCA---TCTTGTAGAACTTgctct | 1nt ins |
| | #2(ε6) | ε | accgccaCCCTGCA-ATCA---TCTTGTAGAACTTgctct<br>accgccaCCCTGCATATCA---TCTTGTAGAACTTgctct | 1nt ins<br>2nt ins |
| CK1δ KO | #1(δ9o.) | δ | gcagctccatca-CCATGA-CGTTGTAGTCCCCCTCTgcccc<br>gcagctccatca-CCATGA-CGTTGTAGTCCCCCTCTgcccc | 2nt del<br>2nt del |
| | #2(δ1) | δ | gcagctccatca-CCATGA-CGTTGTAGTCCCCCTCTgcccc<br>gcagctccatca-CCATGACCGTTGTAGTCCCCCTCTgcccc<br>gcagctccatca-CCATGA-CGTTGTAGTCCCCCTCTgcccc | 1nt del<br>1nt ins<br>11nt del |
| CK1δ/ε* dKO | #1(LH) | ε | accgccaCCCTGC--GGCAAGTTCTTGTAGAACTTgctct<br>accgccaCCCTGCA-ATCA---TCTTGTAGAACTTgctct | 2xSNP + 3nt ins<br>1nt ins |
| | | δ | gcagctccatca-CCATGA-CGGTGTAGTCCCCCTCTgcccc<br>gcagctccatca-CCATGA-CGTTGTAGTCCCCCTCTgcccc | 2nt del + SNP<br>2nt del |

**Figure S3: Production and validation of CK1δ/ε KO panel cell lines.**
**(A)** Sequences of gRNAs used to generate single and compound KO cell lines in the CK1δ/ε KO panel (Fig.1Ei). gRNAs were designed in accordance with PCR screening of KO clones using RFLP analysis. The PAM sequence (in bold) is not part of the gRNA sequence. Underlined sequences indicate recognition/cleavage sites of the specified restriction enzymes (RE). Expected PCR product sizes and corresponding restriction patterns are provided.
**(B)** Agarose gel (2 %) showing RFLP analysis of KO clones.
**(C)** NGS results confirming the KO status of the presented cell lines. PAM sequences are shown in blue and purple for CK1ε and CK1δ isoforms, respectively. gRNA sequences are displayed in black uppercase letters. Multiple sequences within one isoform represent different allele modifications. Dashes are included to improve sequence alignment. ins = insertion; del = deletion; SNP = single nucleotide polymorphism.

### Figure S3

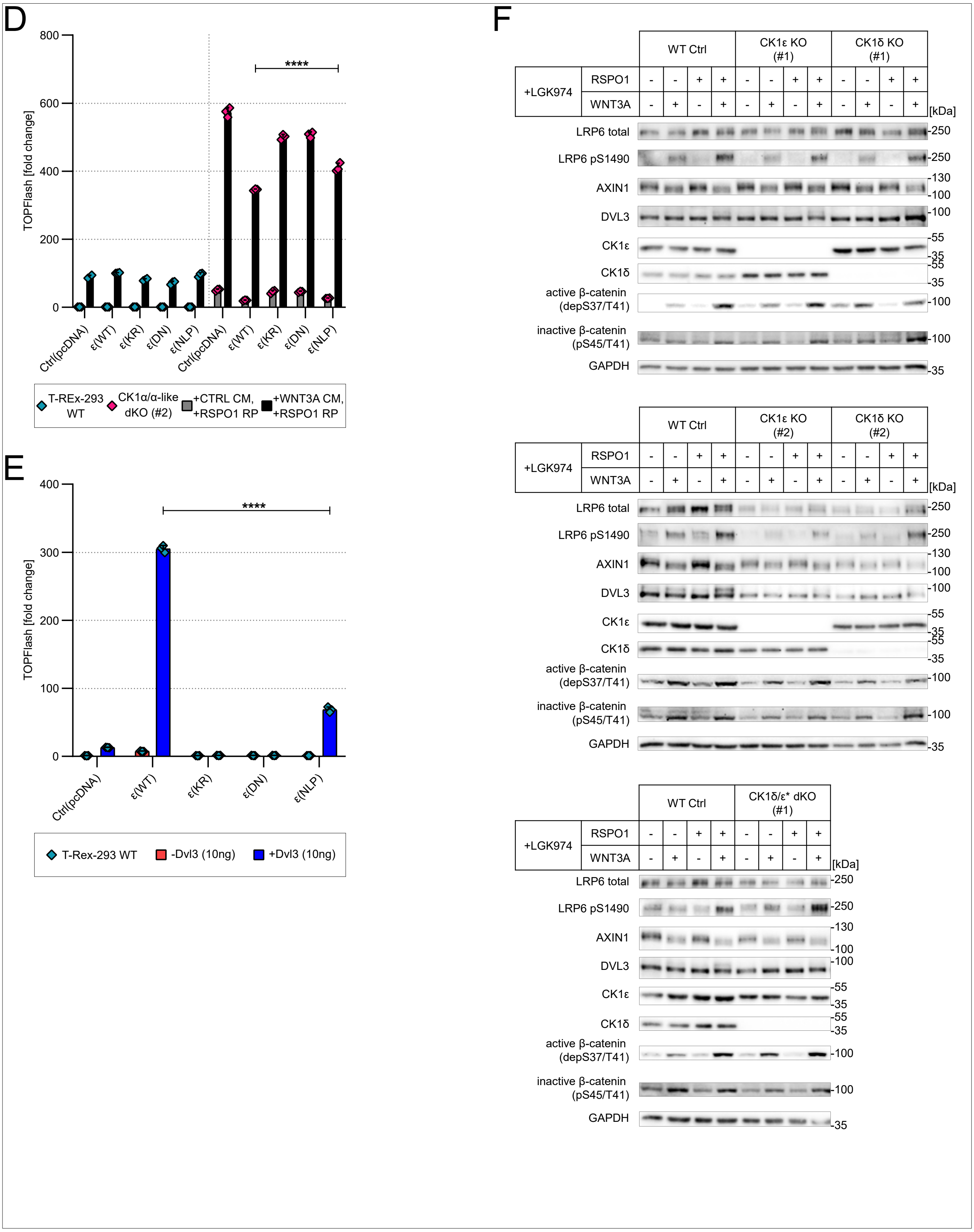


**Figure S3: Production and validation of CK1δ/ε KO panel cell lines.**
**(D)** TOPFlash assay: degradasome assay with CK1ε(WT), kinase-dead mutants, and CK1ε(NLP) mutant.
Statistical analysis: Three-way ANOVA with Tukey's multiple comparisons test (ns [P>0.05], * [P≤0.05], ** [P≤0.01], *** [P≤0.001], **** [P≤0.0001]). Columns represent means with S.D. error bars. n = 3 independent biological replicates.
**(E)** TOPFlash assay: signalosome assay with CK1ε(WT), kinase-dead mutants and CK1ε(NLP) mutant.
Statistical analysis: Ordinary two-way ANOVA with Tukey's multiple comparisons test (ns [P>0.05], * [P≤0.05], ** [P≤0.01], *** [P≤0.001], **** [P≤0.0001]). Columns represent means with S.D. error bars. n = 3 independent biological replicates.
**(F)** Western blot analysis of CK1δ/ε KO panel clones using protein lysates from the TOPFlash analysis (Fig.1Eiii).

### Figure S4

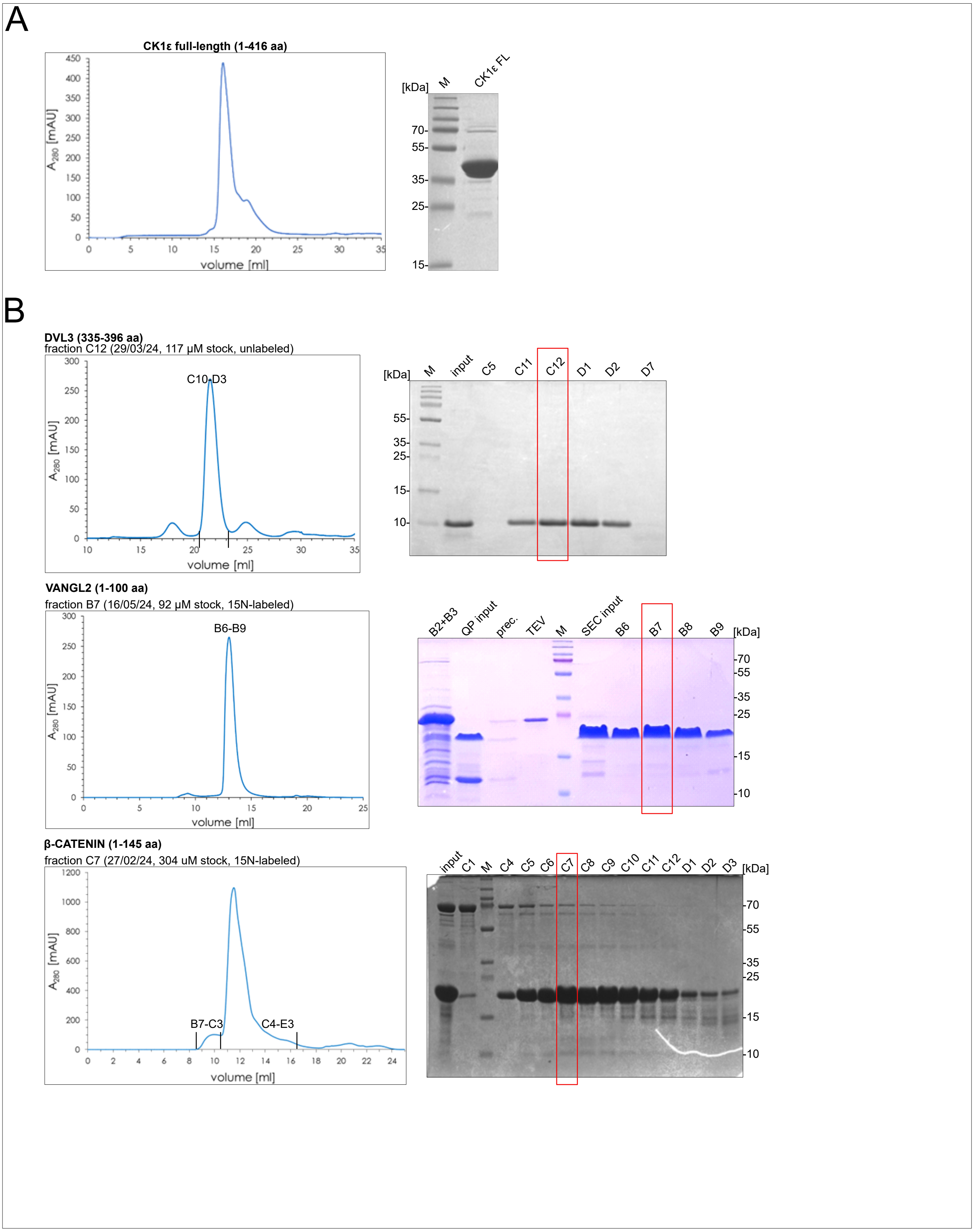


**Figure S4: Purification of substrates and kinases for *in vitro* analyses.**
**(A)** Purification output for kinase CK1ε full-length (1–416 aa). Chromatogram from SEC and SDS-PAGE gel highlighting the fraction used are shown.
**(B)** Purification outputs for substrates DVL3 (335–396 aa), VANGL2 (1–100 aa), and β-catenin (1–145 aa). Chromatograms from SEC and SDS-PAGE gels highlighting the fractions used are shown.

### Figure S5

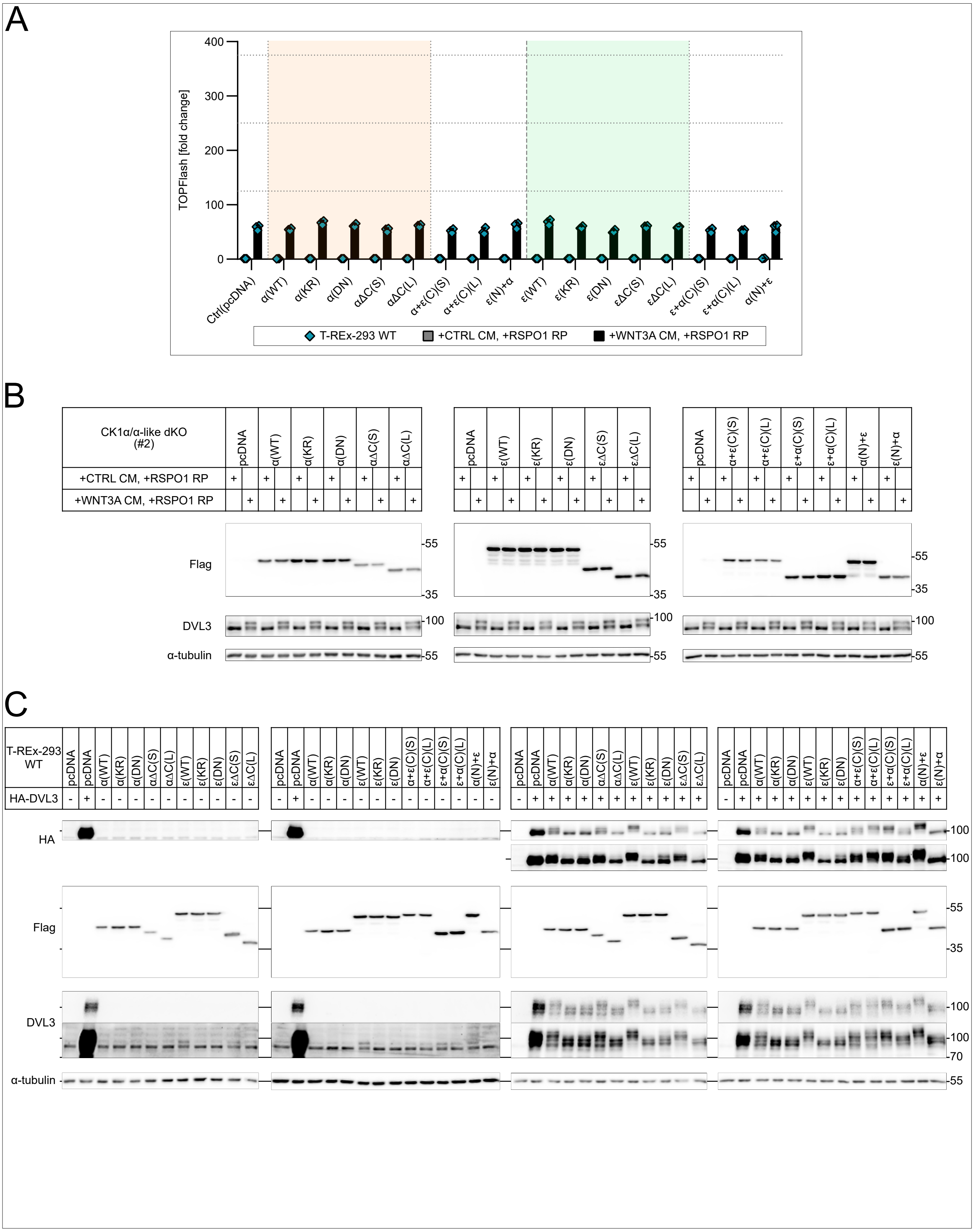


**Figure S5: The N-lobe of CK1α and the C-terminus of CK1ε define cellular preferences for the β-catenin destruction complex and DVL3 signalosome, respectively.**
**(A)** Extension of Fig.3C showing the TOPFlash assay: degradasome assay with N-terminal and C-terminal single swapping mutants between CK1α and CK1ε in T-REx-293 WT cells. Columns represent means with S.D. error bars. n = 3 independent biological replicates.
**(B)** Western blot analysis of TOPFlash lysates from Fig.3C.
**(C)** Western blot analysis of TOPFlash lysates from Fig.3D.

### Figure S5

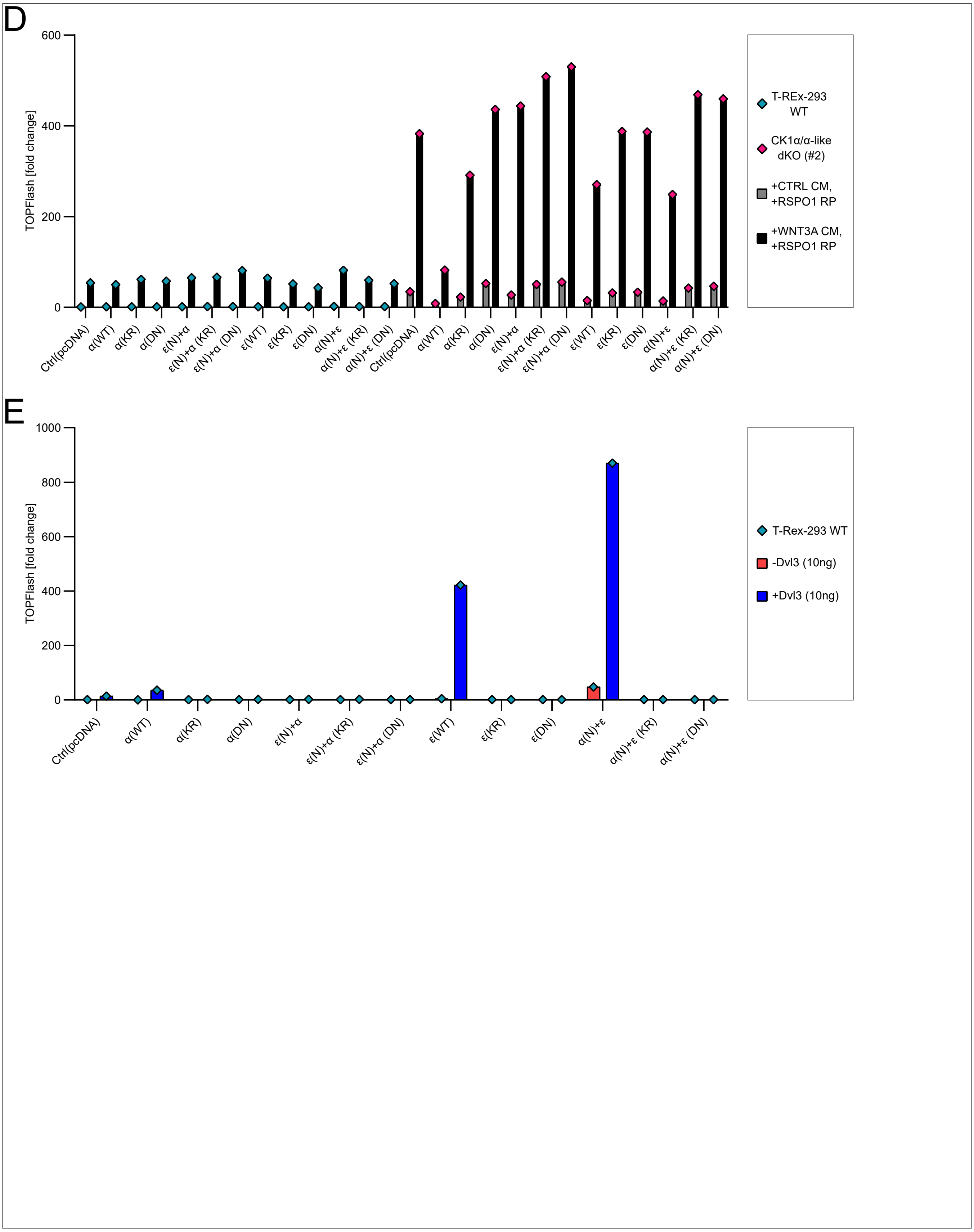


**Figure S5: The N-lobe of CK1α and the C-terminus of CK1ε define cellular preferences for the β-catenin destruction complex and DVL3 signalosome, respectively.**
**(D)** TOPFlash assay: degradasome assay with N-terminal and C-terminal single swapping mutants between CK1α and CK1ε, including their kinase-dead versions. Columns represent means. n = 1 biological replicate.
**(E)** TOPFlash assay: signalosome assay with N-terminal and C-terminal single swapping mutants between CK1α and CK1ε, including their kinase-dead versions. Columns represent means. n = 1 biological replicate.

### Figure S6

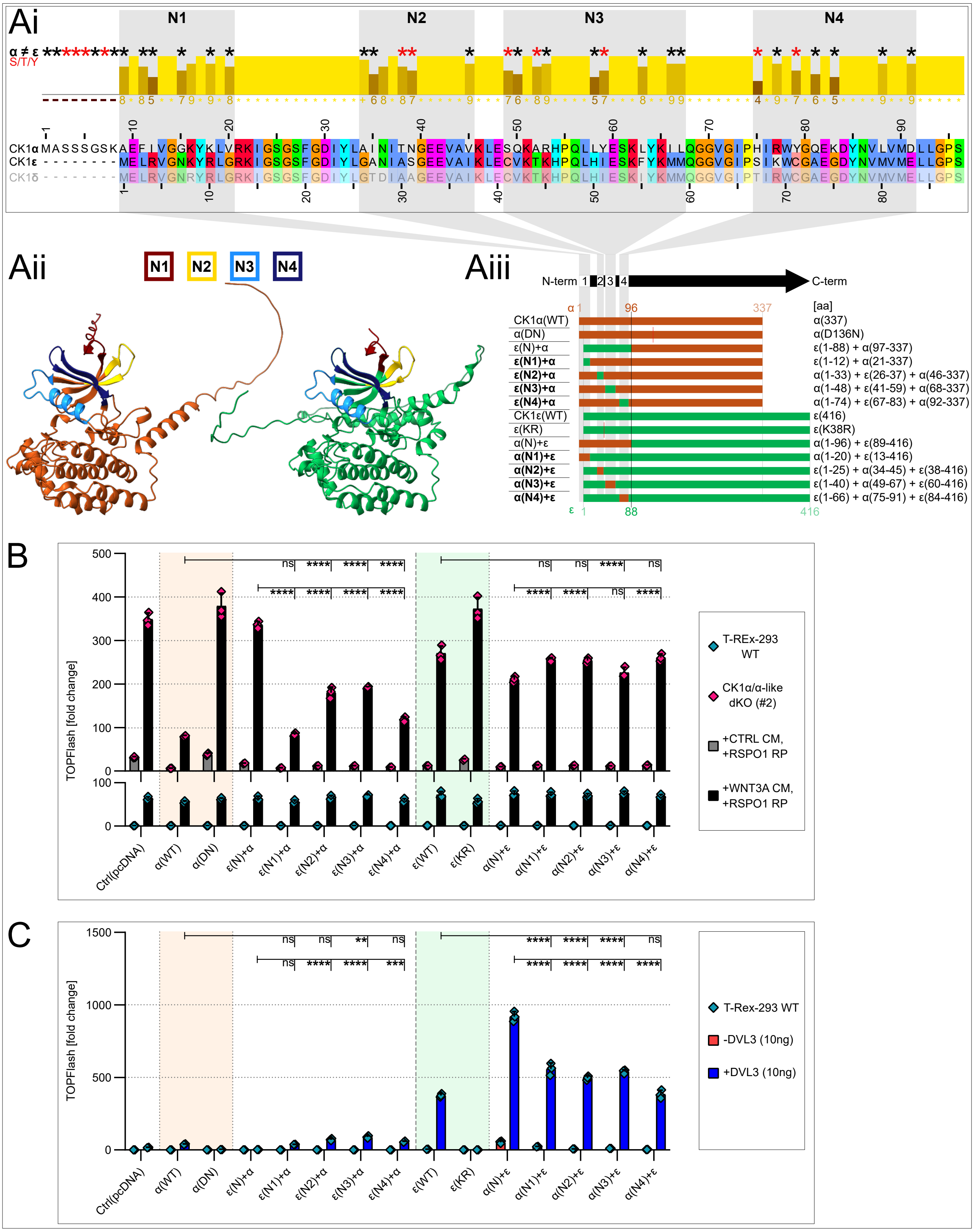


**Figure S6: N-terminal regional swapping mutants suggest the complex nature of identity between CK1α and CK1ε.**
**(A)** Alignment (Ai) of CK1α, CK1ε, and CK1δ highlighting four clusters (N1–N4) with higher residue variability (created in Jalview 2.11.4.1). AlphaFold3 predictions (Aii) depicting N1-N4 regions on the structures of CK1α (orange) and CK1ε (green). Schematic depiction (Aiii) of the panel showing N-terminal regional swapping mutants between CK1α and CK1ε.
**(B)** TOPFlash assay: degradasome assay with mutants from Fig.S6Aiii. Statistical analysis: Three-way ANOVA with Tukey's multiple comparisons test. Highlighted are statistical comparisons for conditions in KO cells.
**(C)** TOPFlash assay: signalosome assay with mutants from Fig.S6Aiii. Statistical analysis: Ordinary two-way ANOVA with Tukey's multiple comparisons test.
**(B,C)** Columns represent means with S.D. error bars. n = 3 independent biological replicates. (ns [P>0.05], * [P≤0.05], ** [P≤0.01], *** [P≤0.001], **** [P≤0.0001]).

### Figure S6

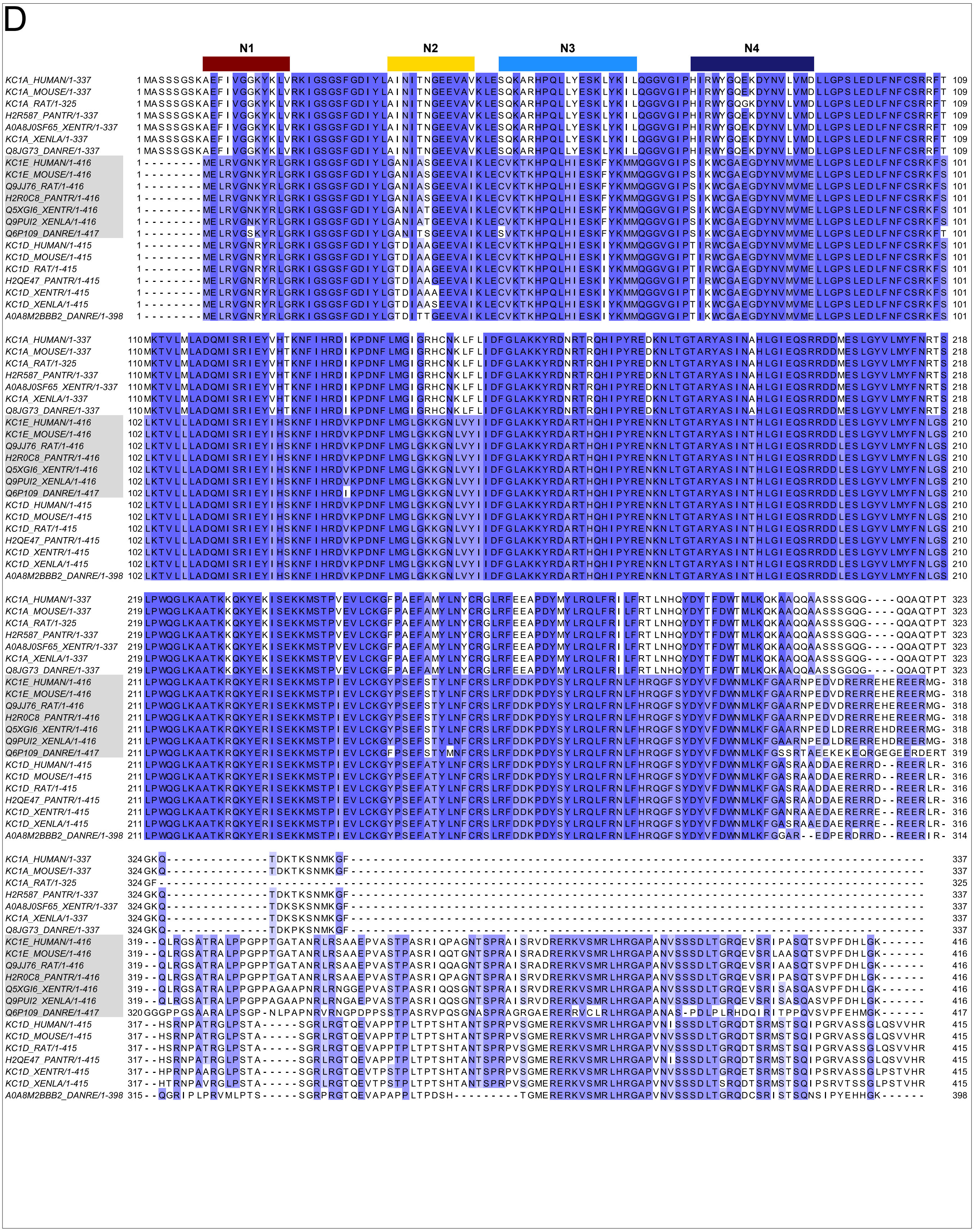


**Figure S6: N-terminal regional swapping mutants suggest the complex nature of identity between CK1α and CK1ε.**
**(D)** Evolutionary alignment of CK1α, CK1δ, and CK1ε from *Homo sapiens, Mus musculus, Rattus norvegicus, Pan troglodytes, Xenopus tropicalis, Xenopus laevis,* and *Danio rerio*. Highlighted are the N1-N4 regions from Fig.S6Aii. The alignment was created in Jalview 2.11.4.1. Coloring is based on percentage identity.

### Figure S7

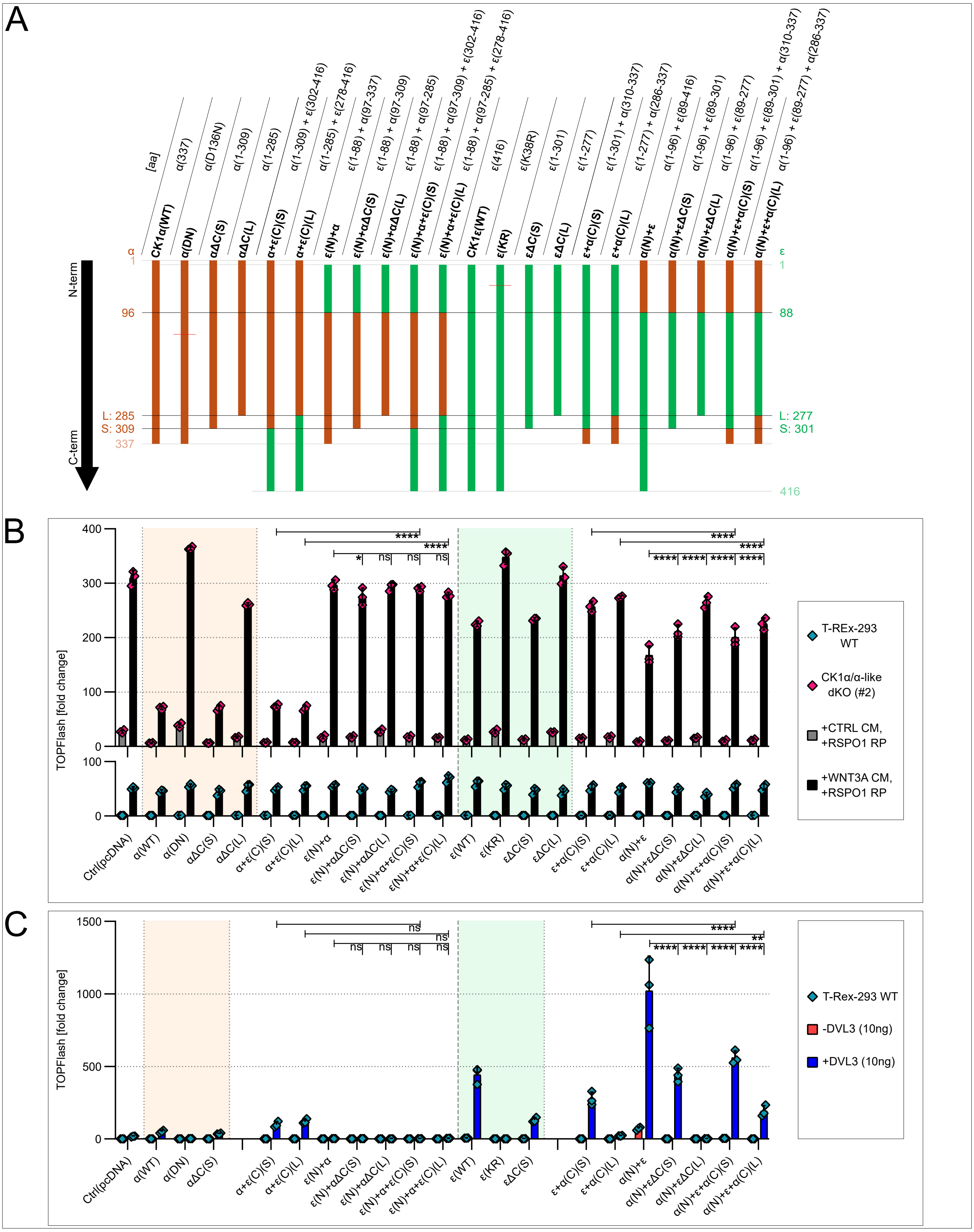

**Figure S7: Double swapping mutants exclude C-terminal interference with N-lobe function.**
**(A)** Schematic depiction of the panel showing N-terminal and C-terminal double swapping mutants between CK1α and CK1ε.
**(B)** TOPFlash assay: degradasome assay with mutants from Fig.S7A. Statistical analysis: Three-way ANOVA with Tukey's multiple comparisons test. Highlighted are statistical comparisons for conditions in KO cells.
**(C)** TOPFlash assay: signalosome assay with mutants from Fig.S7A. Statistical analysis: Ordinary two-way ANOVA with Tukey's multiple comparisons test.
**(B,C)** Columns represent means with S.D. error bars. n = 3 independent biological replicates. (ns [P>0.05], * [P≤0.05], ** [P≤0.01], *** [P≤0.001], **** [P≤0.0001]).

### Figure S8

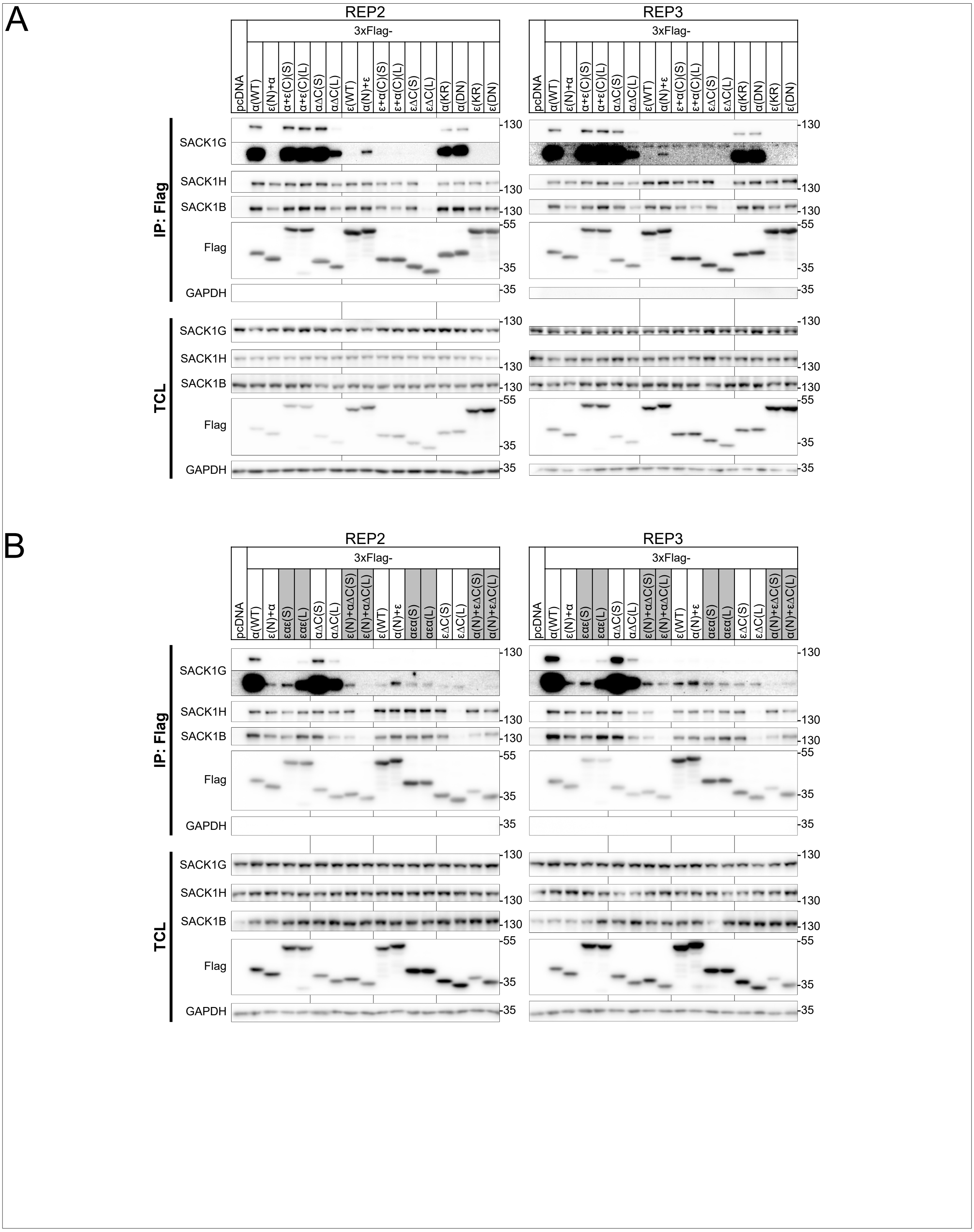


**Figure S8: The N-terminal region of CK1α is a structural determinant of its selective interaction with SACK1G (FAM83G).**
**(A)** Additional replicates to Fig.4B of co-immunoprecipitation: Immunoprecipitation of 3xFlag-tagged CK1 isoforms, their mutants, and chimeric constructs (panel from Fig.3B) in overexpression conditions, with detection of pulled-down SACK1G, SACK1H, and SACK1B. TCL - total cell lysate input.
**(B)** Additional replicates to Fig.4C of co-immunoprecipitation: Immunoprecipitation of 3xFlag-tagged CK1 isoforms, their mutants, and chimeric constructs (panel from Fig.S7A) in overexpression conditions, with detection of pulled-down SACK1G, SACK1H, and SACK1B. TCL - total cell lysate input.

### Figure S8

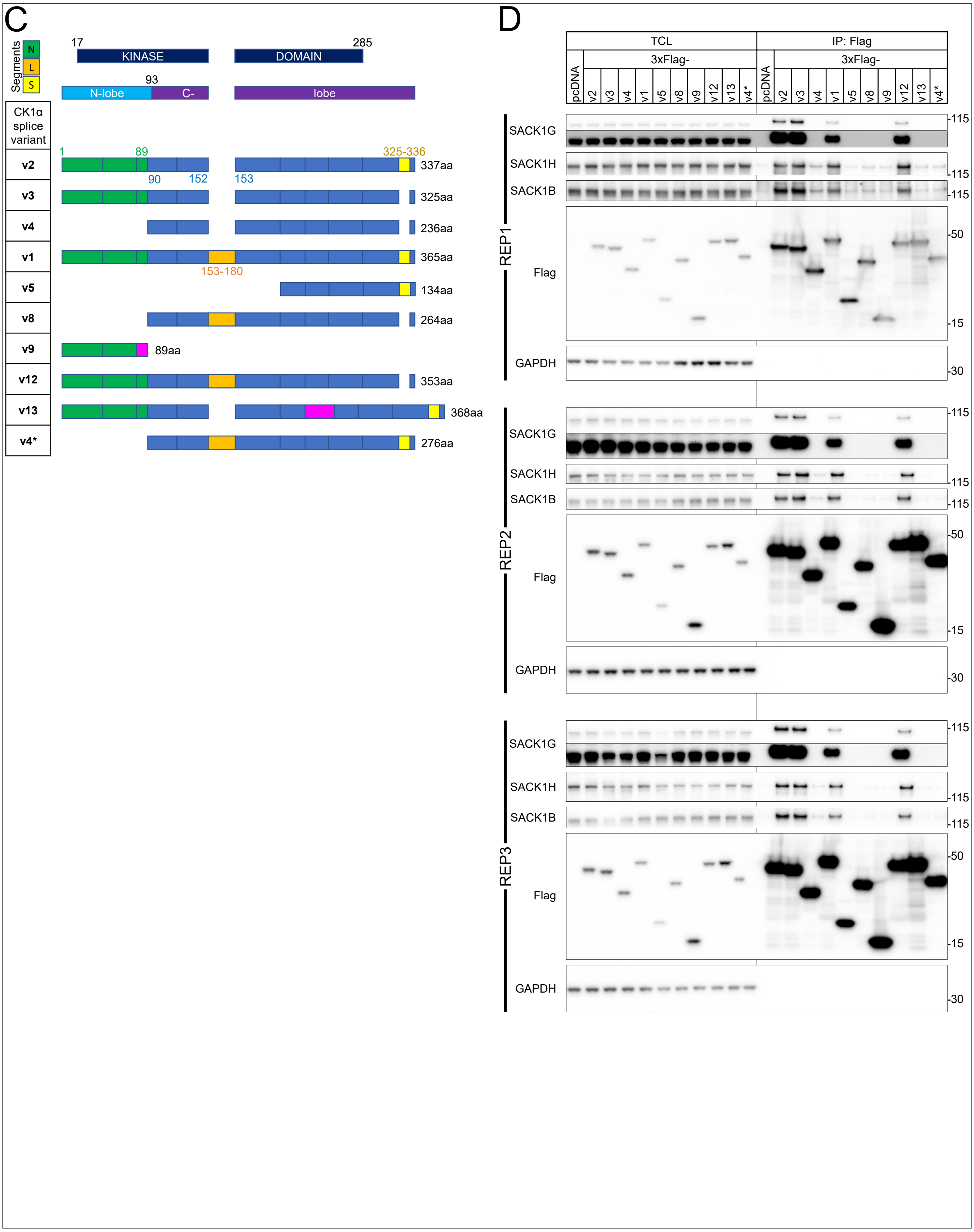


**Figure S8: The N-terminal region of CK1α is a structural determinant of its selective interaction with SACK1G (FAM83G).**
**(C)** Schematic depiction of CK1α splice variants. Adapted from Gybel *et al.*, 2024.
**(D)** Co-immunoprecipitation: Immunoprecipitation of 3xFlag-tagged CK1α splice variants from Fig.S8C in overexpression conditions, with detection of pulled-down SACK1G, SACK1H, and SACK1B. Three biological replicates are presented. TCL - total cell lysate input.

### Figure S9

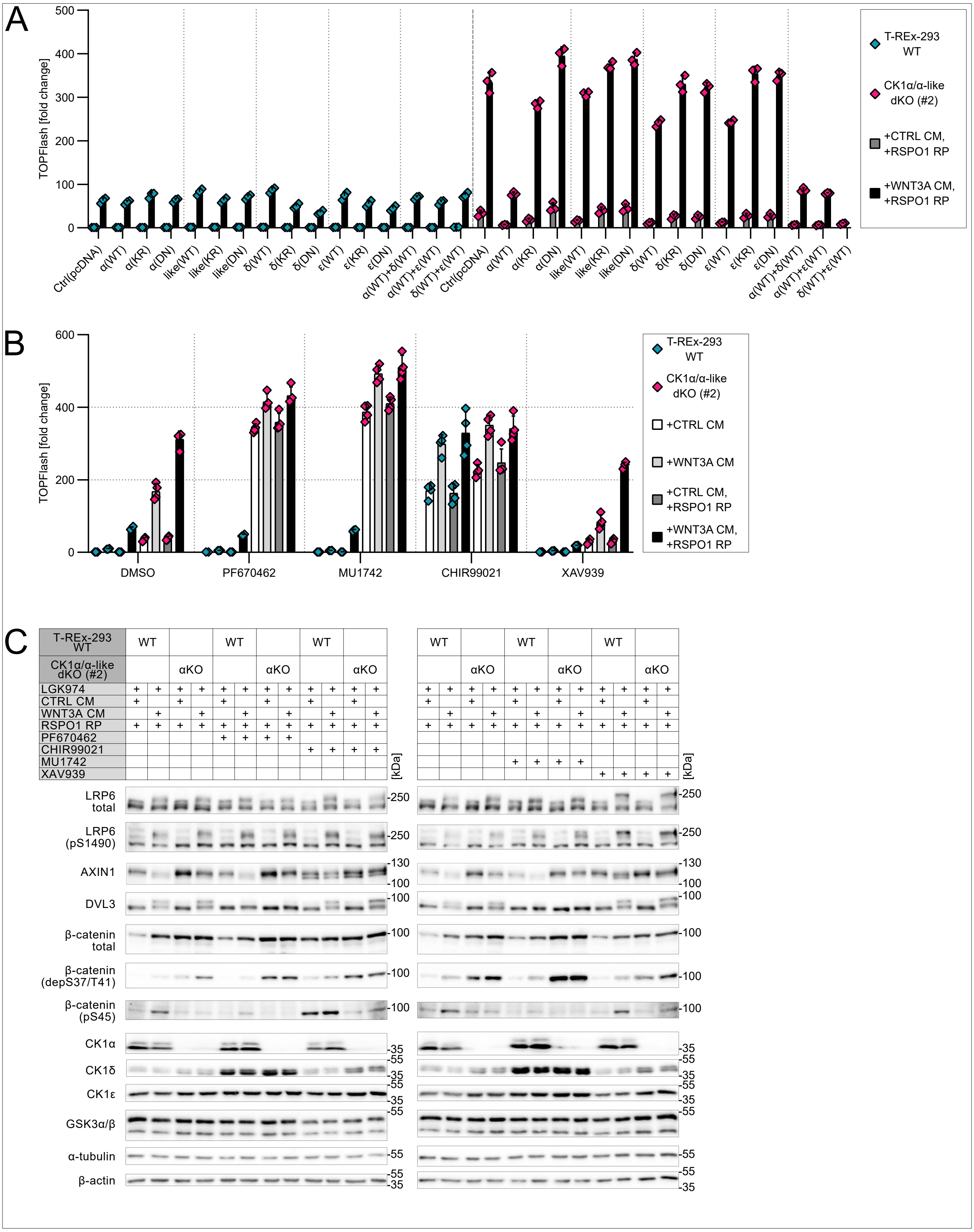


**Figure S9: In the absence of CK1α, CK1δ and CK1ε physically and functionally replace CK1α in the destruction complex.**
**(A)** Complete data set from Fig.5A, including kinase-dead mutants and measurements in T-REx-293 WT cells.
**(B)** Complete data set from Fig.5B, including treatments with CTRL CM and WNT3A CM without RSPO1 RP.
**(C)** Western blot analysis of TOPFlash lysates from Fig.5B and Fig.S9B.

### Figure S9

D

| Next Generation Sequencing Results | | | | |
|---|---|---|---|---|
| CSNK1A1 gene - hg19: chr5 (+) 148892662 – 148892694 | | | | |
| WT | | α | gctagCATATCGGGCAGTGCC-AGTGAGGttttt | |
| KO lines | | | | Type of modification |
| CK1α/δ dKO | #1(A2) | δ | based on CK1δ KO (#1) in Fig.S3C | |
| | | α | gctagCATATCGGGCAGTGCCAAGTGAGGttttt | 1nt ins |
| | #2(C3) | δ | based on CK1δ KO (#1) in Fig.S3C | |
| | | α | gctagCATATCGGGCAGTGCCAAGTGAGGttttt | 1nt ins |
| CK1α/ε dKO | #1(A2) | ε | based on CK1ε KO (#1) in Fig.S3C | |
| | | α | gctagCATATCGGGCAGTGCCAAGTGAGGttttt | 1nt ins |
| | #2(D4) | ε | based on CK1ε KO (#1) in Fig.S3C | |
| | | α | gctagCATATCGGGCAGTGCCAAGTGAGGttttt | 1nt ins |
| CK1α/δ/ε* tripleKO | #1(3.1) | δ/ε* | based on CK1δ/ε* dKO (#1) in Fig.S3C | |
| | | α | gctagCATATCGGGCAGTGCC-AGTGAGGttttt | 10nt del |
| | #2(4.1) | δ/ε* | based on CK1δ/ε* dKO (#1) in Fig.S3C | |
| | | α | gctagCATATCGGGCAGTGCCAAGTGAGGttttt | 1nt ins |

E

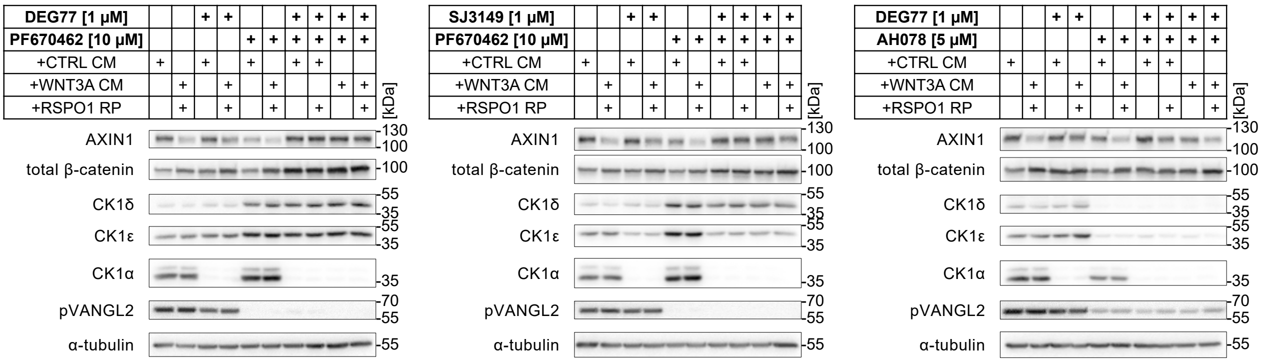


F

| Next Generation Sequencing Results | | | | |
|---|---|---|---|---|
| CSNK1A1 gene - hg19: chr5 (+) 148892662 – 148892694 | | | | |
| WT | | α | gctagCATATCGGGCAGTGCC-AGTGAGGttttt | |
| KO lines | | | | Type of modification |
| CK1α + DVL1/2/3 tetraKO | #1(F1) | α | gctagCATATCGGGCAGTGCCAAGTGAGGttttt | 1nt ins |
| | | DVL1/2/3 | based on DVL1/2/3 tripleKO in Paclíková *et al.*, 2017 | |
| | #2(E1) | α | gctagCATATCGGGCAGTGCCAAGTGAGGttttt | 1nt ins |
| | | DVL1/2/3 | based on DVL1/2/3 tripleKO in Paclíková *et al.*, 2017 | |
| CK1α + LRP5/6 tripleKO | #1(C6) | α | gctagCATATCGGGCAGTGCCAAGTGAGGttttt | 1nt ins |
| | | LRP5/6 | based on LRP5/6 dKO in Bernatik *et al.*, 2021 | |
| | #2(A5) | α | gctagCATATCGGGCAGTGCCAAGTGAGGttttt | 1nt ins |
| | | LRP5/6 | based on LRP5/6 dKO in Bernatik *et al.*, 2021 | |

G

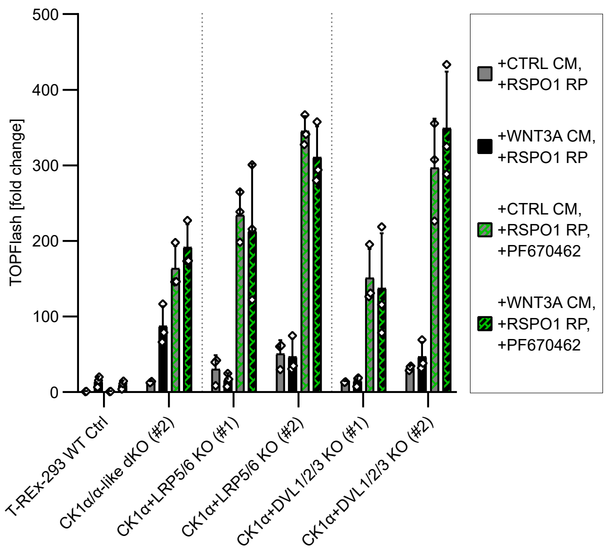


**Figure S9: In the absence of CK1α, CK1δ and CK1ε physically and functionally replace CK1α in the destruction complex.**
**(D)** NGS results confirming the KO status of the presented cell lines. The PAM sequence is shown in blue. The gRNA sequence is displayed in black uppercase letters. Dashes are included to improve sequence alignment. ins = insertion; del = deletion.
**(E)** Western blot analysis of TOPFlash lysates from Fig.5D.
**(F)** NGS results confirming the KO status of the presented cell lines. The PAM sequence is shown in blue. The gRNA sequence is displayed in black uppercase letters. Dashes are included to improve sequence alignment. ins = insertion.
**(G)** TOPFlash assay testing genetic models of CK1α depletion in the background of LRP5/6 dKO and DVL1/2/3 tripleKO. Cells were cultured in the absence or presence of PF670462 (10 μM for 16 h) and Wnt/β-catenin pathway stimulation along with 0.5 μM LGK974. Columns represent means with S.D. error bars. n = 3 independent biological replicates.
